# Modeling lung adenocarcinoma using layer-by-layer nanoparticles mitigates innate immune cell activation

**DOI:** 10.64898/2026.09.14.751530

**Authors:** Vidit Bhandarkar, Elen Torres-Mejia, Taylor A. Heim, Namita Nabar, Tamara G. Dacoba, Zachary J. Rogers, Nicolas Mathey-Andrews, Sean-Luc Shanahan, Chau Vo, Adam Berger, J. Christopher Love, Tyler Jacks, Paula T. Hammond, Stefani Spranger

**Affiliations:** Department of Biology at MIT; Koch Institute for Integrative Cancer Research at MIT; Department of Chemical Engineering at MIT; 4Harvard Medical School; Program in Health Sciences and Technology at Harvard and MIT; Broad Institute of MIT and Harvard; Institute for Soldier Nanotechnologies; Ragon Institute of MGH, MIT and Harvard

**Author notes:** These authors contributed equally (Vidit Bhandarkar, Elen Torres-Mejia, Taylor A. Heim).

## Abstract

Lung adenocarcinoma, driven frequently by KRAS and p53 mutations, remains a leading cause of cancer mortality. Current state-of-the-art genetically engineered mouse models often rely on viral delivery of recombinases, such as Cre recombinase, to initiate transformation. However, viral particles can infect and activate innate immune cells, thus potentially impacting studies of tumor-immune dynamics. Here, we develop a layer-by-layer (LbL) polyplex platform using poly(β-aminoester) (PBAE) polymers layered with poly-L-aspartic acid (PLD) to deliver Cre mRNA to lungs while avoiding immune cell transfection and activation. PLD-coated nanoparticles (PLD-NPs) exhibit stable mRNA encapsulation and efficient transfection in vitro, even after lyophilization and long-term storage. In Kras^LSL-G12D/+^;p53^flox/flox^ (KP) mice, PLD-NPs initiate lung adenocarcinomas that mirror human histopathology without infecting or activating dendritic cells and alveolar macrophages, unlike lentiviral (LV) or adenoviral delivery methods. Single-cell transcriptional profiling revealed that LV administration drives long-term upregulation of antigen presentation and costimulatory machinery in lung-resident myeloid populations. This persistent immune activation is avoided by NP delivery. By uncoupling tumor initiation from innate immune activation, this platform enables the high-fidelity interrogation of tumor-immune dynamics, especially for non-inflammation-driven lung cancer.

## Introduction

Lung cancer remains the leading cause of cancer-related mortality worldwide^1^, with non-small cell lung cancer (NSCLC) accounting for 85-90% of all cases^1, 2^. Lung adenocarcinoma (LUAD) is the most common histological subtype of NSCLC, and KRAS mutations are among the most frequent oncogenic drivers in this context, detected in over one third of lung adenocarcinomas^3–7^. These mutations, particularly at codon 12 (such as G12C, G12D, and G12V), are early and ubiquitous events in tumor evolution and are associated with high tumor mutation burden and immune evasion phenotypes ^6, 7^. Immune checkpoint blockade (ICB) therapies have transformed the treatment landscape for NSCLC and emerged as standard of care for a subset of patients with Kras-mutant tumors^8, 9^. However, a significant fraction of patients fail to respond to treatment, highlighting the urgent need for improved preclinical models to better understand disease mechanisms and immune evasion, especially in non-inflammation-induced lung cancer.

Our mechanistic understanding of the immune response to lung cancer is tightly linked to the development of preclinical mouse models^10–13^. Syngeneic transplantable models of lung cancer are extensively utilized because they are rapid and amenable to genetic and pharmacological perturbations. However, these models fail to recapitulate natural lung tumor progression, a key aspect of patient pathophysiology, making it challenging to dissect the dynamic interactions between the immune system and evolving tumors. In contrast, inducible genetically engineered mouse models (GEMMs) of lung cancer faithfully mimic tumor progression, enabling interrogation of the complex interactions between developing tumors and the immune system.

Kras-driven LUAD is commonly studied using the well-established Cre recombinase-inducible KP (Kras^lox-stop-lox(LSL)-G12D/+^; p53^flox/flox^) GEMM^14–17^. Cre is often delivered intratracheally via adenoviral (Ad) or lentiviral (LV) vectors which induce transient or sustained Cre expression respectively. This model accurately recapitulates several features of human lung cancer, including tumor progression and immune evasion^14, 17^. However viral delivery of Cre to initiate transformation introduces several challenges in modeling immune dynamics including (1) lack of cell type specificity in viral transduction leading to viral vector delivery to non-target cells (2) expression of viral transgenes by these cells and (3) innate immune sensing of the viral particle itself. Specifically, the use of ubiquitous promoters such as phosphoglycerate kinase (PGK-LV) promoter, lack cell-type specificity, and can result in infection of immune cells, including antigen-presenting cells (APCs) such as dendritic cells (DC) and alveolar macrophages (AM)^10, 18–20^. In contrast, tissue-specific promoters such as the surfactant C promoter (SPC) or Clara cell secretory protein promoter (CCSP), more tightly restrict Cre expression to lung epithelial cells. However, intratracheal Ad-SPC transduces more CD45+ leukocytes than Epcam+ epithelial cells in the lung, demonstrating that tissue-specific promoters do not completely circumvent expression outside of the desired cell type ^10^. Furthermore, the use of tissue-specific promoters may restrict expression of delivered transgenes, but it does not prevent viral entry and integration into non-target cells.

The use of LV for Cre delivery leads to sustained Cre expression which carries its own set of hurdles. Cre recombinase causes toxicity in a dose-dependent manner as well as chromosomal abnormalities ^21, 22^. Furthermore, the sustained expression of a foreign protein (Cre) is a potential neo-antigen within itself, though it should be noted that there are no reports identifying antigen-specific T cell responses against Cre. The NINJA (iNversion-INducible Joined neoAntigen) model represents an elegant method to decouple transformation and antigen expression by using a two-step Flp and Cre-dependent inversion cassette ^23^. This two-step process prevents neoantigen expression in the thymus, thereby avoiding central tolerance. However, because the NINJA model relies on viral delivery of Cre recombinase to the lung, there is still the potential for off-target transgene integration and for viral particles to infect and activate myeloid cells, thereby affecting the tumor microenvironment.

The potential for transducing immune cells when using viral Cre delivery is not a novel concern, and several methods have been used in recent years to overcome this issue. Transduction of lung immune cells can be minimized by reducing the total amount of virus and by incorporating microRNA target sequences into the expression cassette that are complementary to microRNAs in hematopoietic cells ^24–26^. While these methods effectively and drastically minimize transduction of innate immune cells, it is unclear whether they completely prevent innate immune cell sensing of viral vectors and ensuing activation.

Numerous studies have demonstrated that viral infection in the lungs can reprogram the innate immune system, enhancing its responsiveness to unrelated secondary stimuli ^27–29^. One alternative to viral induction are spontaneous latent alleles such as the Kras^LA1^ GEMM mouse model, which relies on global spontaneous recombination. While this model is not dependent on viral delivery of recombinases, it is subject to onset of tumor formation in organs other than the lung, making it challenging to study immune dynamics in the tumor ^30^. These limitations of viral-based delivery of Cre, or the reliance on spontaneous transformation underscores the need for alternative approaches, specifically if the designed studies focus on myeloid cells within the tumor microenvironment.

To overcome these challenges, we turned to synthetic nanoparticles (NPs), which can enable targeted and efficient delivery of nucleic acids. Polyplexes, or polymeric nanosystems, can stably complex nucleic acids into nanoscale structures through electrostatic interactions. Furthermore, NP surface chemistries can be engineered for targeting or stealth advantages using Layer-by-layer (LbL), or the electrostatic adsorption of charged outer coatings. LbL polyplexes offer modular assembly, tunable surface properties, and enhanced protection of mRNA cargo ^31–34^. By engineering polyplexes to evade uptake by and prevent activation of innate immune cells, it is possible to achieve high-fidelity genetic manipulation in vivo. Such specificity is particularly valuable for modeling oncogenic events, such as Cre-mediated recombination in conditional lung cancer models, without confounding effects from immune cell transfection or activation.

Here, we developed a NP platform using poly(β-aminoester) (PBAE)-based polyplexes layered with the bioactive polypeptide poly-L-aspartic acid (PLD) to deliver Cre mRNA to lung epithelial cells, while minimizing uptake by innate immune cells such as DCs and AMs. We used this system to initiate LUAD in KP mice and observed tumor progression consistent with tumors induced by PGK-LV vectors. Notably, transient Cre expression via this platform led to efficient tumor induction without transfection or activation of professional APCs. Transcriptional profiling revealed increased activation of dendritic cells and alveolar macrophages in PGK-LV-induced tumors 14 weeks after administration with observed consequences for the activation state of tumor-infiltrating T cells. This approach expands the existing toolkit for modeling NSCLC by enabling precise genetic manipulation with minimal inflammation and immune perturbation at the time of tumor initiation.

### Intratracheal delivery of viral vectors shapes local and systemic immune responses

To assess the impact of PGK-LV infection on innate immune cells within the lung, we intratracheally (I.T.) injected TdTomato^LSL/LSL^ (T) reporter mice with a phosphoglycerate kinase promoter-driven lentiviral vector encoding Cre recombinase, which activates TdTomato expression in transfected cells. (Fig. 1A). We have chosen the PGK-LV platform due to its broad use and all studies below use this platform unless specified otherwise. Flow cytometric analysis performed 7 days later revealed that a significant fraction of DCs and AM were infected by PGK-LV (Fig. 1B, Supp. Fig. 1A-C).

**Figure 1:**
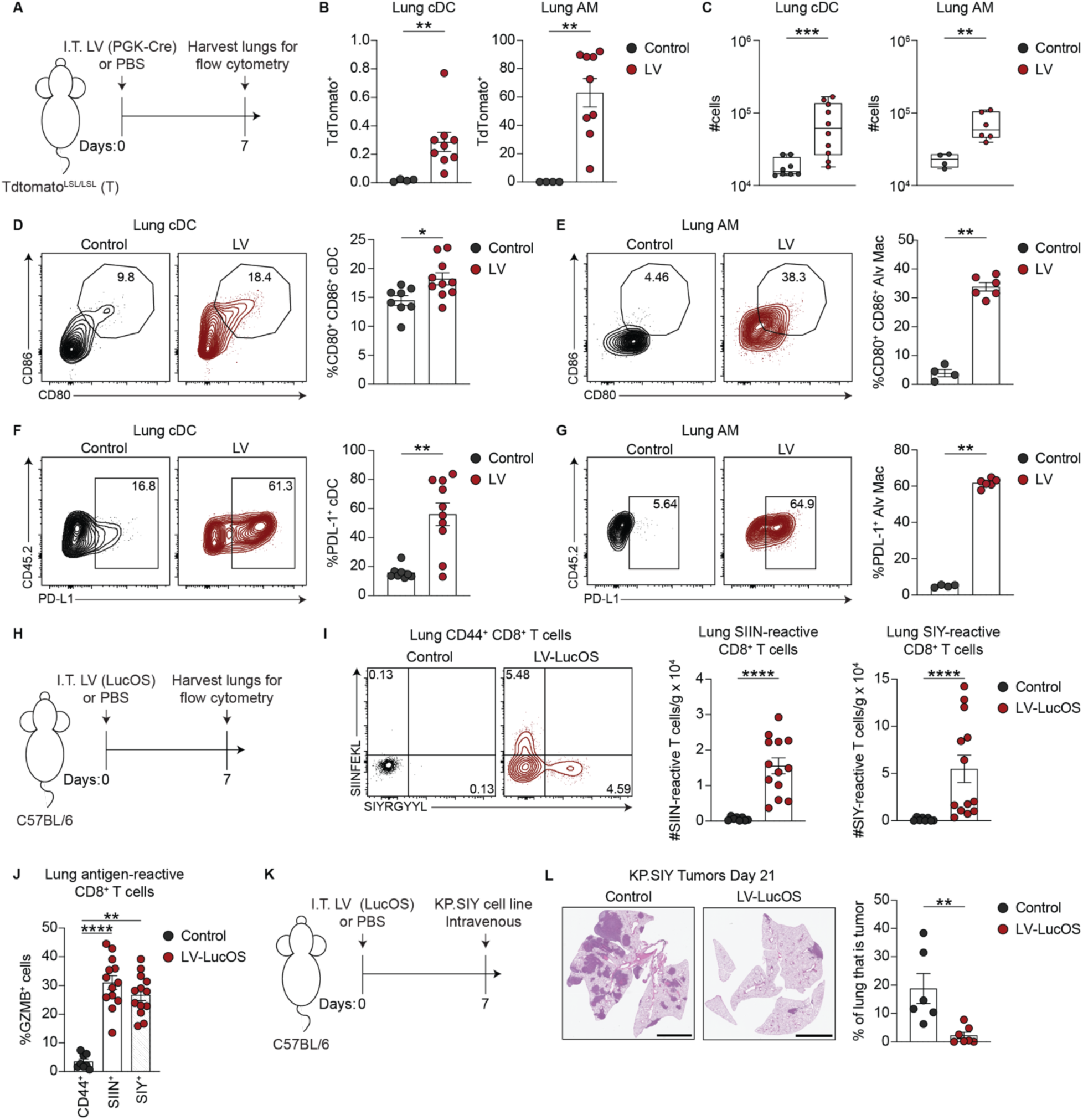
Intratracheal delivery of lentiviruses elicits a robust immune response. (A) Experimental scheme: intratracheal (I.T.) administration of PGK-LV to TdTomato^LSL/LSL^ reporter mice. (B-G) Analyses of lung dendritic cell (DC) and alveolar macrophage (AM) populations. (B) Percent of TdTomato-positive DCs and AM (Control n=4, LV n=9). (C) Total count of lung DCs (DC control n=8, DC LV n=10) and AMs (AM control n=4, AM LV n=6). (D, E) Expression of CD80 and CD86 on lung DCs and AMs (n=8-10 DCs, n=4-6 AMs). (F, G) PD-L1 on lung DCs and AMs, respectively (n=8-10 DCs, n=4-6 AMs). (H) Experimental scheme for I and J. LucOS LV (encoding Cre, SIINFEKL, and SIYRYYGL). (I) Quantification and representative flow plots of lung SIIN- and SIY-reactive CD8+ T cells in LucOS-infected mice (control n=8, LucOS n=13). (J) Percentage of GZMB-positive SIIN- or SIY-reactive CD8+ T cells in the lung (control n=8, LucOS n=13). (K) Experimental scheme for LucOS priming followed by I.V. challenge with KP.SIY tumor cells. (L) Quantification and representative H&E images of lung tumor burden 21 days after KP.SIY challenge (control n=6, LucOS n=7). Scale bar = 4000μm. Mann-Whitney U test used for statistical calculations in B-G, I, L; one-way ANOVA for J. Data are presented as mean ± SEM.

Next, we investigated whether PGK-LV administration activates APCs in the lungs and draining mediastinal lymph nodes (mLN) in C57BL/6 mice. PGK-LV administration increased the number of DCs (Fig. 1C, Supp. Fig. 1D) as well as the expression of costimulatory molecules CD40, CD80, and CD86 on DCs in the lungs (Fig. 1D Supp. Fig. 1E) and mLN (Supp. Fig. 1F-H), indicating strong activation. AMs in the lungs also increased in number and exhibited elevated CD80 and CD86 expression (Fig. 1C, 1E). Furthermore, PD-L1 was upregulated on both DCs and AMs (Fig. 1F-G), indicative of an inflammatory response likely driven by IFNγ^35^. Consistent with previous reports^10^, I.T. injection of an adenoviral vector encoding Cre under the lung epithelial-specific surfactant protein C (SPC) promoter (Ad-SPC-Cre) also led to off-target immune cell activation (Supp. Fig. 1I-L). These data demonstrate that lenti and adenoviral particles broadly infect and activate myeloid cells in the lung and draining LN following intratracheal delivery.

To evaluate the functional consequences of APC infection and activation, we utilized the LucOS PGK-LV ^14, 36, 37^, which encodes Cre along with two CD8^+^ T cell antigens SIINFEKL and SIYRYYGL that bind MHC-I H2-Kb. Infecting C57BL/6 mice with LucOS PGK-LV (Fig. 1H) allows us to track antigen-specific CD8^+^ T cell responses independent of tumor formation (Supp. Fig. 2A). We observed a striking increase in the number of CD44^+^ CD8^+^ T cells in the lungs and mLN of mice 7 days post I.T. injection (Supp. Fig. 2B-C), indicative of robust T cell activation. This was accompanied by a significant expansion of antigen-specific CD8^+^ T cell populations (Fig. 1I, Supp. Fig. 2D), which expressed activation markers such as CD25 and granzyme B (GZMB) (Fig. 1J, Supp. Fig. 2E-F). These findings indicate that LucOS PGK-LV administration, despite being a tool for tumor initiation, can induce a strong T cell response even in the absence of tumors, potentially influencing tumor outgrowth and confounding interpretations of immune dynamics in LV-based models.

**Figure 2.**
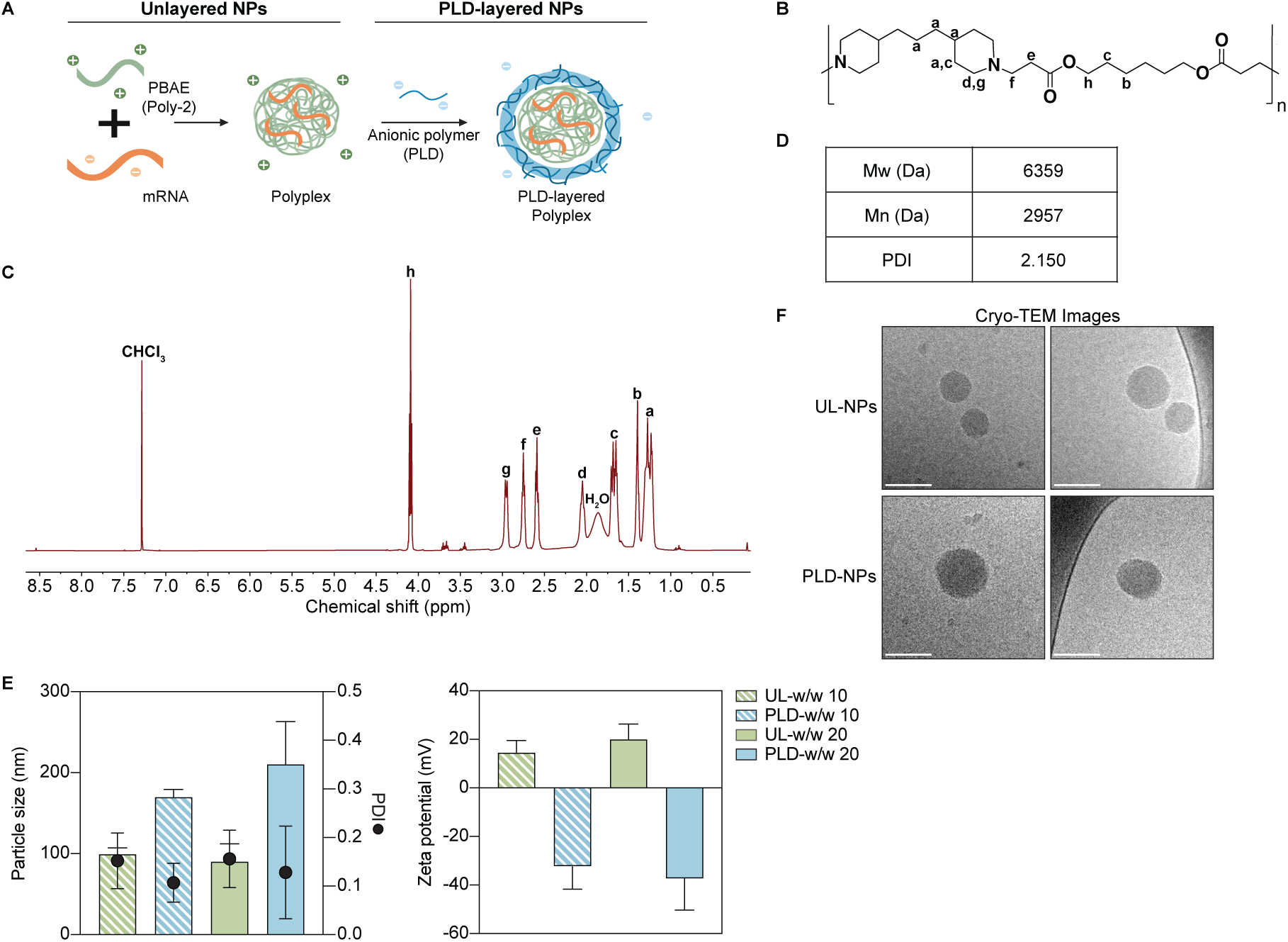
mRNA-polyplexes can be electrostatically layered with bioactive polyanions. A) Schematic illustrating polyplex formation through electrostatic interaction between poly(β-amino esters) (PBAEs) and mRNA, followed by layering with anionic polymers. B) Chemical structure of Poly2, the PBAE used for polyplex formulation. C) Representative 1D ¹H NMR spectrum of Poly2 in deuterated chloroform (CDCl₃), with characteristic peaks labeled. D) Weight-average molecular weight (Mw), number-average molecular weight (Mn), and polydispersity index (PDI) of synthesized Poly2, determined by gel permeation chromatography. E) Hydrodynamic diameter, PDI, and zeta potential of UL- and PLD-NPs formulated at Poly2:mRNA weight ratios of 10 and 20. For w/w 10: N=3–4 batch replicates with 3 technical replicates each. For w/w 20, UL-NPs: n=76 batch replicates, 1-3 technical replicates each. PLD-NPs: n=28 batch replicates, 1-3 technical replicates each. Data shown as mean ± SD. F) Representative cryo-transmission electron microscopy images of UL-NPs and PLD-NPs; scale bar is 100nm.

To directly test whether this premature immune activation alters tumor progression, we challenged LucOS-infected C57BL/6 mice with LUAD tumor cells expressing the model antigen SIYRYYGL (called KP.SIY) via tail vein injection (Fig. 1K). Mice that had previously received LucOS PGK-LV exhibited a marked reduction in lung tumor burden when rechallenged 1 week after initial infection (Fig. 1L). Notably, this effect persisted in a memory setting; delaying tumor rechallenge by four weeks yielded the same protective outcome (Supp. Fig. 3A-B). Given that activated, antigen-reactive CD8^+^ T cells were detected in the spleen and blood post-PGK-LV infection (Supp. Fig. 3C-E), we assessed whether localized LucOS PGK-LV administration in the lungs could confer systemic protection against subcutaneous tumors. Indeed, when mice were challenged with KP.SIY flank tumors at both seven days and five weeks post-LucOS infection, we observed delayed tumor outgrowth (Supp. Fig. 3F-I), affirming that LucOS PGK-LV administration induces systemic anti-tumor immunity.

**Figure 3:**
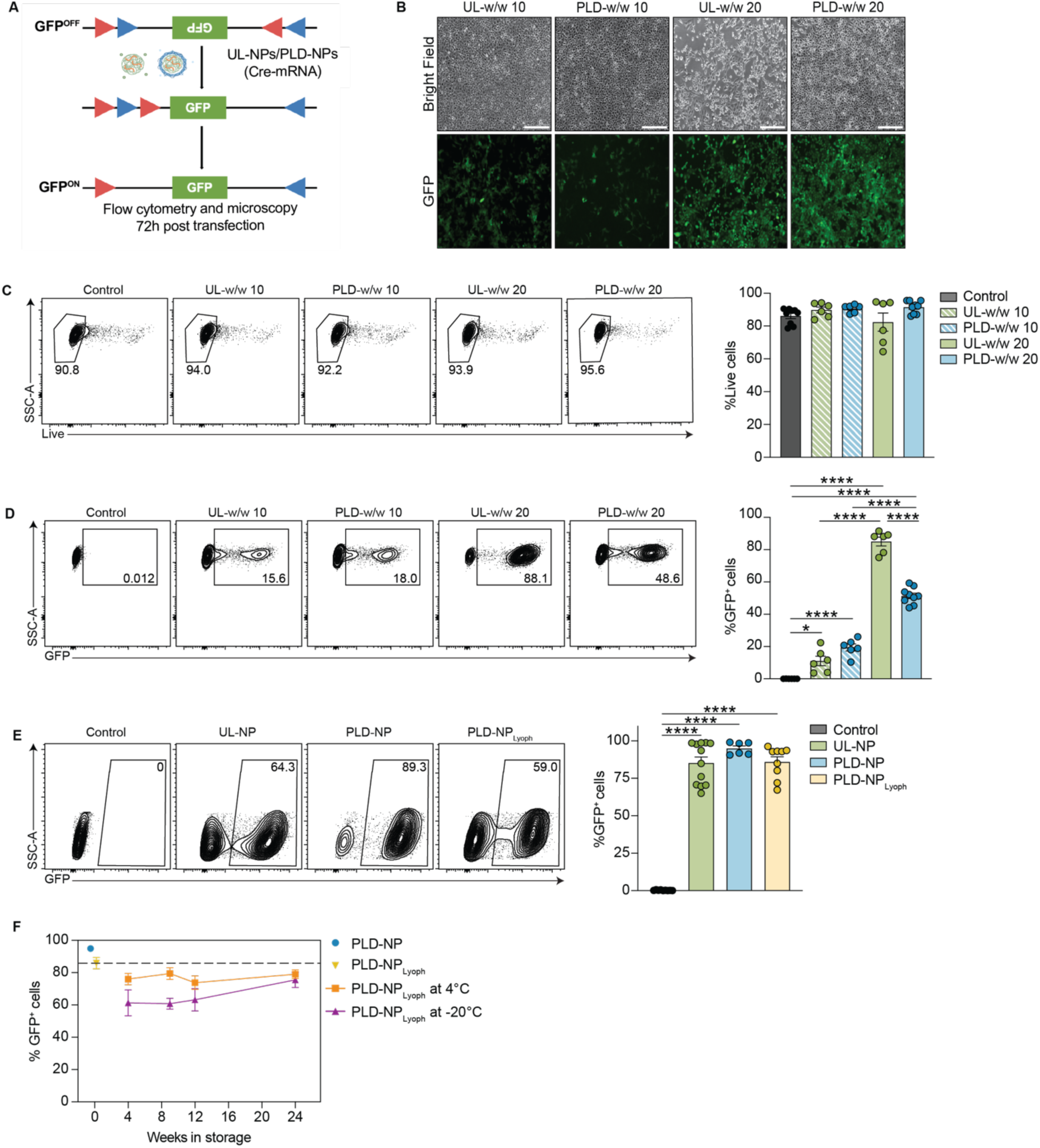
PLD-NPs demonstrate high transfection and retain potency upon lyophilization. A) Schematic depicting the Green-Go Cre-reporter cell line response to delivery of Cre mRNA-containing NPs (100 ng mRNA per well). 72 h after treatment with UL- or PLD-NPs, cells were analyzed by flow cytometry or microscopy. B) Representative bright-field and fluorescence microscopy images of Green-Go cells treated with UL- or PLD-NPs at w/w 10 or 20, n=4. Scale bar = 250μm. C) Representative flow plots (left) and quantification (right) of live cells after treatment with UL- or PLD-NPs at w/w 10 or 20, n=6. D) Representative flow plots and percentage of GFP^+^ cells following transfection with UL- or PLD-NPs at w/w 10 or 20, n=6. E) Representative flow plots and quantification of GFP^+^ cells following treatment with UL-NPs, PLD-NPs, or PLD-NPLyoph, n=6-12. F) Percent GFP^+^ cells following treatment with freshly prepared PLD-NPs or PLD-NPLyoph stored at 4°C or –20°C for up to 24 weeks. n=6-12. For C-F, P-values calculated using one-way ANOVA and data shown as mean ± SD.

### mRNA-polyplexes can be electrostatically layered with bioactive polyanions

Seeking a delivery strategy that minimally alters immune cells, we turned to non-viral, nanoparticle-based delivery systems. With sizes similar to viruses, their composition can be engineered to effectively deliver genetic cargos to cells of interest^38, 39^. Both lipid-based nanoparticles (Lipid-NPs) and polymer-based nanoparticles (polyplexes) have shown robust nucleic acid transfection capacity. Nevertheless, prior studies have reported stronger immune responses against lipid-NPs compared to polyplexes in vaccine applications^40^. Here, we leveraged the high transfection capacity of cationic polymers such as PBAEs to electrostatically complex nucleic acids into polyplexes (Fig. 2A). For improved stability and reduced toxicity, we further layered the resulting polyplexes with PLD, a negatively charged polyaminoacid.

Among PBAE designs, Poly2 has demonstrated potent nucleic acid delivery in multiple contexts, including pDNA delivery via polyplexes and siRNA release from electrostatic thin films^32–34, 41^. Here, we examined the ability of this polymer to encapsulate and deliver mRNA in polyplexes. Poly2 was synthesized via stepwise polymerization (Fig. 2B)^32^. Polymer structure was confirmed via 1D and 2D 1H NMR (Fig. 2C, Supp. Fig. 4A), and molecular weight was measured as Mw = 6359 Da via gel permeation chromatography (Fig. 2D, Supp. Fig. 4B). Unlayered polyplexes (UL-NP) composed of Poly2 and Cre-mRNA, were first formed and then layered with an outer coating of PLD (PLD-NP) (Fig. 2A).

**Figure 4.**
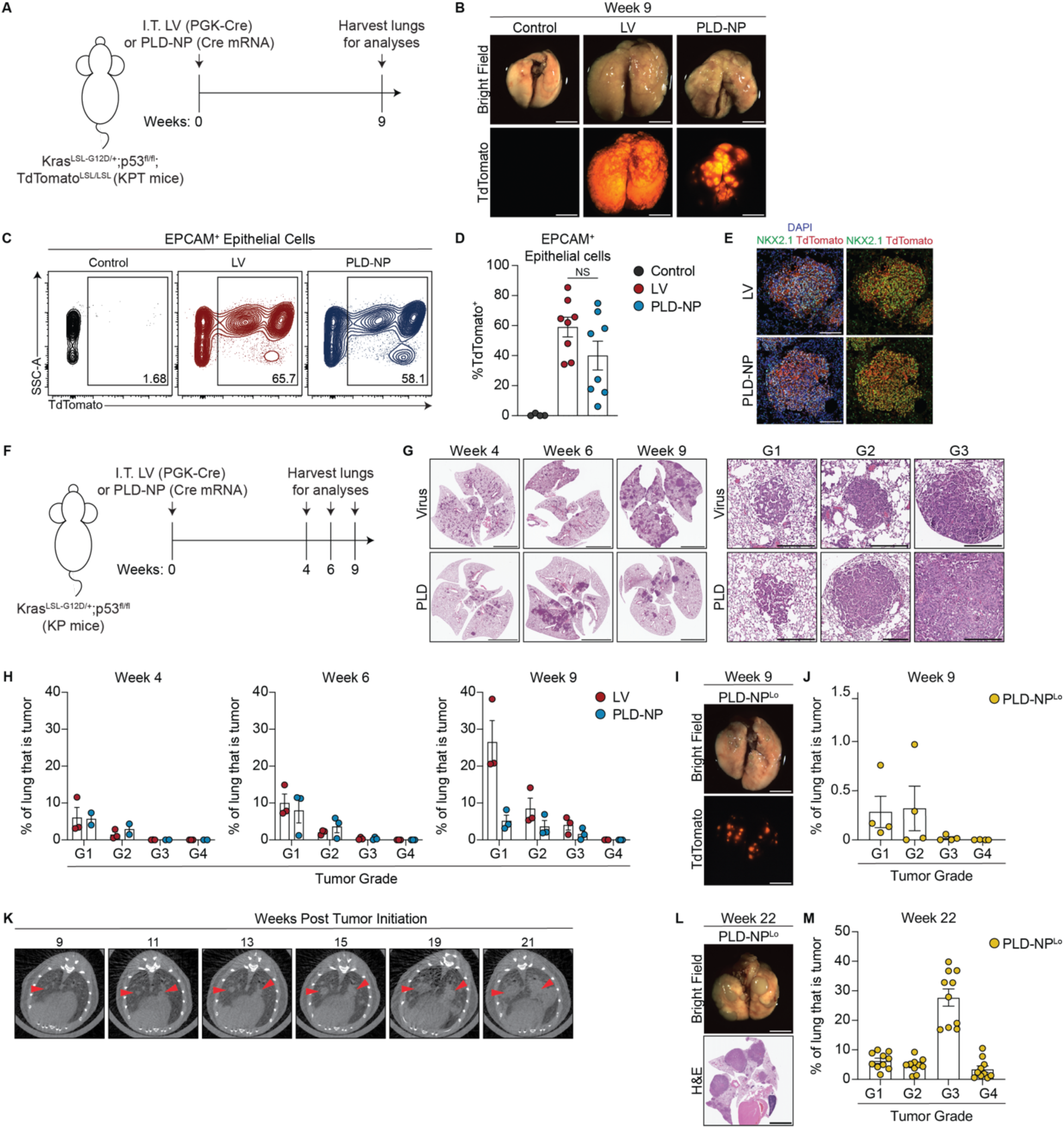
Intratracheal delivery of Cre-mRNA encapsulating PLD-NPs initiates LUAD and recapitulates human disease progression and histopathology. A) Experimental scheme. B) Bright-field and fluorescent images of lungs at week 9, showing TdTomato. Scale bar. = 500 pixels. (C-D) Representative flow cytometry plots (C) and quantification (D) of TdTomato^+^ EPCAM^+^ epithelial cells 9 weeks after tumor initiation, control n=4, LV n=9, PLD-NP n=7, P-values calculated using Mann-Whitney U test. E) Immunofluorescence of lung sections showing colocalization of NKX2.1 and TdTomato 9 weeks after tumor induction. F) Experimental schematic for (G-H). G) Representative H&E-stained lung sections, showing tumor burden and histologic grade. Scale bar = 4000μm (left), or 300μm (right). H) Tumor burden by grade (G1–G4) at weeks 4, 6, and 9, LV n= 3, PLD n=2-3. I) Brightfield and fluorescent images of lungs at week 9 post I.T. injection of PLD-NP^lo^. Scale bar = 500 pixels. J) Quantification of tumor burden by grade in PLD-NP^lo^–injected mice at week 9, n=4. K) Representative μCT scans of KP mice injected with PLD-NP^lo^. Lung tumors marked with red arrows. L) Representative whole lung (scale bar = 500 pixels) and H&E images (scale bar = 4000μm) from PLD-NP^lo^ injected mice at endpoint (week 22). M) Quantification of tumor burden by histologic grade at week 22; n=10. All data shown as mean ± SEM.

To evaluate the impact of polymer quantity on NP formation and transfection, we generated polyplexes at two weight ratios of polymer to mRNA, 10 and 20, of polymer to mRNA. Via dynamic light scattering (DLS), both formulations showed hydrodynamic diameters of 90-100 nm, polydispersity indices (PDI) below 0.3 suggesting relatively homogeneous populations, and positive surface zeta potentials of +15 and +20 mV, respectively (Fig. 2E). Upon layering with PLD, both formulations showed a size increase of 70 – 100 nm, and complete reversal of charge to ∼ -40 mV (Fig. 2E). The modularity of the LbL assembly enabled using alternative bioactive and bioinert polymers for polyplex layering, such as poly-L-glutamic acid (PLE), hyaluronic acid (HA), or poly(acrylic acid). We observed high mRNA encapsulation in all formulations, as measured by gel electrophoresis and RiboGreen quantification of intact polyplexes (Supp. Fig. 4D-E). Via cryo-transmission electron microscopy, polyplexes demonstrated comparable, electron-dense morphologies that appeared undisturbed upon layering (Fig. 2F), further suggesting that layered polyplexes retain stability and mRNA encapsulation. Notably, of the four polyanions tested, PLD maintained polyplex stability at all tested quantities, though PLE and PAA exhibited comparable stable layering. Thus, we proceeded to evaluate PLD-NPs but the incorporation of alternate surface polymers varying in chemistry could further expand the applications of this non-viral delivery tool to other cells and tissues of interest.

### PLD-NPs demonstrate high transfection in vitro and retain potency upon lyophilization

To evaluate the transfection efficiency of both UL- and PLD-NPs at multiple Poly2:mRNA weight ratios, we utilized 3TZ Green-Go reporter cells, which are genetically engineered to express GFP upon Cre exposure (Fig. 3A)^12^. 72h after dosing NPs encapsulating Cre mRNA, cells were evaluated for transfection via flow cytometry and microscopy. We only tested weight ratios of Poly2:mRNA of 10 and 20, given that NPs with a Poly2:mRNA weight ratio of 30 induced high levels cytotoxicity (Supp. Fig. 5A), precluding their further use. The other two NP formulations maintained high cell viability, though UL-NPs with a Poly2:mRNA weight ratio of 20 induced a slight decrease in viability, seen both via microscopy (Fig. 3B) and viability staining (Fig. 3C). Notably, higher Poly2:mRNA weight ratios significantly improved transfection; both UL- and PLD-NPs formulated with a Poly2:mRNA weight ratio of 20 outperformed corresponding groups at a weight ratio of 10 (Fig. 3D). Though PLD-NPs transfected at a lower efficiency compared to UL-NPs, both conditions generated significant GFP signal compared to control (Fig. 3D). Cationic NPs are known to cause cytotoxicity in vitro and in vivo, partially due to potential disruptions of cell membranes^42^. Therefore, the anionic outer layer of PLD serves to mitigate toxic side effects while maintaining sufficient transfection potency. Based on this characterization, we opted to advance the UL- and PLD-NPs with weight ratio of 20.

**Figure 5.**
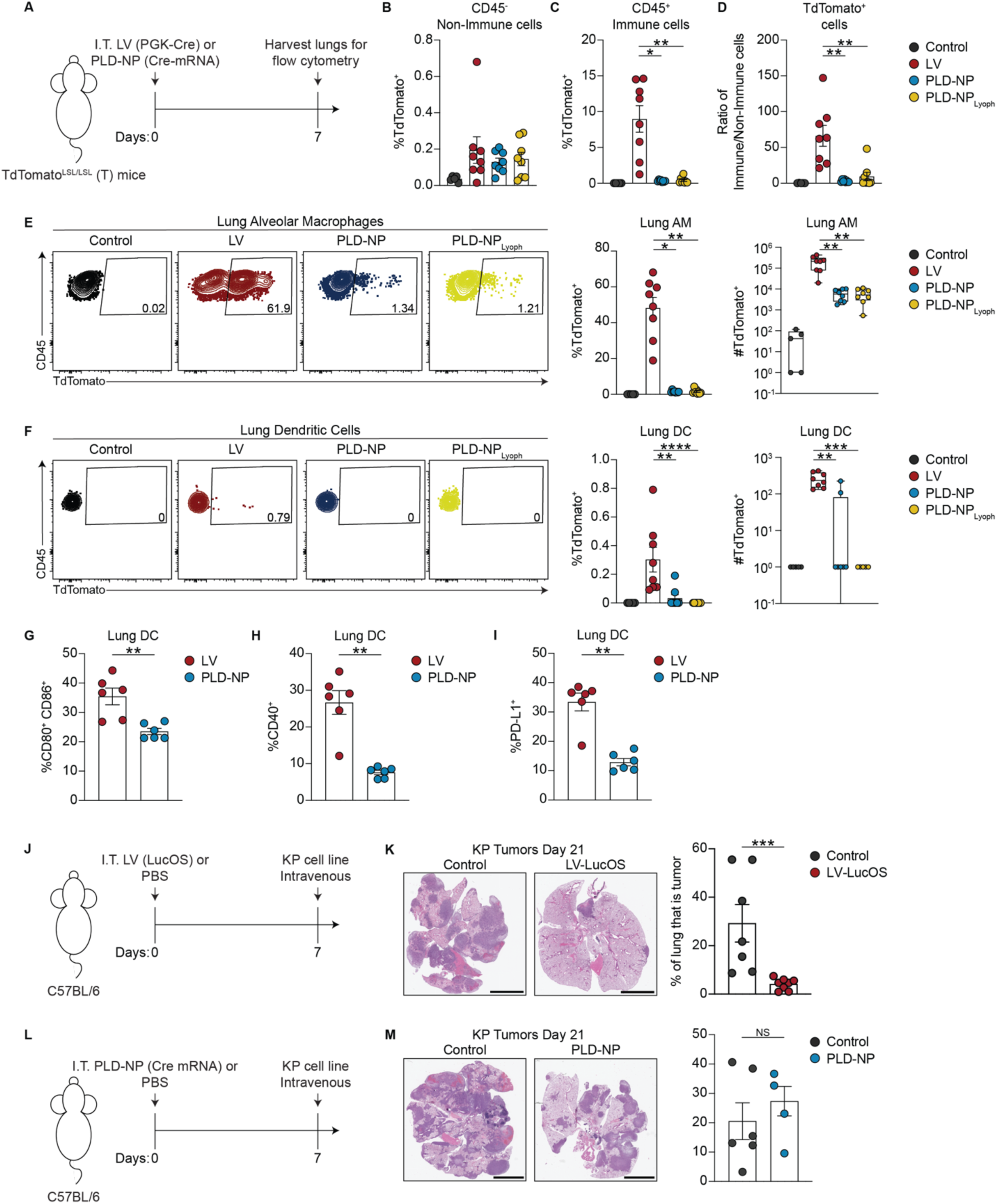
PLD-NPs achieve epithelial cell-specific delivery of Cre-mRNA without innate immune transfection or activation. A) Experimental scheme: TdTomato^LSL/LSL^ (T) reporter mice were I.T. injected with PGK-Cre LV, PLD-NPs, or PLD-NP_Lyoph_ encapsulating Cre-mRNA. Lungs were harvested 7 days later for flow cytometry. B) Percentage of TdTomato^+^ cells among CD45^-^ non-immune cells. C) Percentage of TdTomato^+^ cells among CD45^+^ immune cells. D) Ratio of the frequency of TdTomato^+^ immune to non-immune cells in lungs. E-F) Representative flow plots (left) and quantification (right) of TdTomato^+^ AM (E) and DCs (F), shown as percent and number. For B-F, Control n=5, LV n=8, PLD-NP n=8, PLD-NP_Lyoph_ n=8. G-I) Percentage of lung DCs expressing CD80 and CD86 (G), CD40 (H), and PD-L1 (I). n=6 per group. J) experimental scheme showing I.T. LucOS priming followed by I.V. challenge with parental KP cells lacking SIIN and SIY antigens. (K) Quantification and representative H&E images of lung tumor burden 21 days after challenge with parental tumor cells n=5-8. Scale bar = 4000μm. (L-M) Same as J-K but with PLD-NP instead of LV. Scale bar = 4000∝μ. For B-F quantification, P-values calculated using one-way ANOVA. For G-I, and K, M quantification, P-values calculated using Mann-Whitney U test. All data shown as mean ± SEM.

PBAE polyplexes present limited long-term stability due to the susceptibility of the polymer backbone to hydrolysis. However, lyophilization, or freeze-drying, is a valuable, commercially-employed approach to improve NP shelf life, storage, and handling^43^. Thus, we optimized the polyplex lyophilization process to enable long-term storage. First, we decreased the salt concentration to adsorb PLD, improving NP physical stability over time (Supp. Fig. 5B); consequently, the optimized PLD-NPs demonstrated improved transfection efficiency (Supp. Fig. 5C). Next, we compared lyophilization with two widely used cryoprotectants, sucrose and trehalose at different concentrations. Using the optimized PLD-NPs, we observed an increase in NP size upon lyophilization, irrespective of the sugar and concentration used (Supp. Fig. 5D). 5% sucrose was the lowest concentration able to maintain stable PLD-NPs in isotonic conditions (Supp. Fig. 5D). Therefore, we proceeded with 5% sucrose as our lyophilization strategy and further characterized these NPs (PLD-NP_Lyoph_). As expected, PLD-NP_Lyoph_ maintained a PDI less than 0.3, a negative charge, and the same mRNA encapsulation efficiency following lyophilization (Supp. Fig. 5E-G). We next confirmed that PLD-NP_Lyoph_ could efficiently transfect 3TZ Green-Go reporter cells in vitro despite the changes in particle size (Fig. 3E). Importantly, lyophilization enabled extended storage of PLD-NPs with minimal physicochemical degradation for up to 6 months at both 4°C and -20°C. However, 4°C proved to be the optimal storage temperature for maintaining transfection efficiency. Over a 9-week period, transfection efficiency decreased by only 7.5% at 4°C compared to a 29.2% drop at -20°C (Fig. 3F, Supp. Fig. 5H). Ultimately, this optimization establishes PLD-NP_Lyoph_ as a highly stable delivery platform.

### Intratracheal delivery of PLD-NPs encapsulating Cre-mRNA induces LUAD in KP GEMM that recapitulates human disease progression and histopathology

To achieve sufficient Cre mRNA for in vivo dosing, both UL- and PLD-NPs were concentrated, without significantly altering their physical properties and transfection efficiency in vitro (Supp. Fig. 6A-B). However, UL-NPs exhibited marked in vivo toxicity, inducing significant weight loss and 40% mortality within one week of administration (Supp. Fig. 6C-D). In surviving mice, lungs were abnormally enlarged by day 7 compared to untreated animals and those treated with PGK-LV or PLD-NPs (Supp. Fig. 6E-F). Based on these findings, we proceeded with PLD-NPs for subsequent experiments.

**Figure 6:**
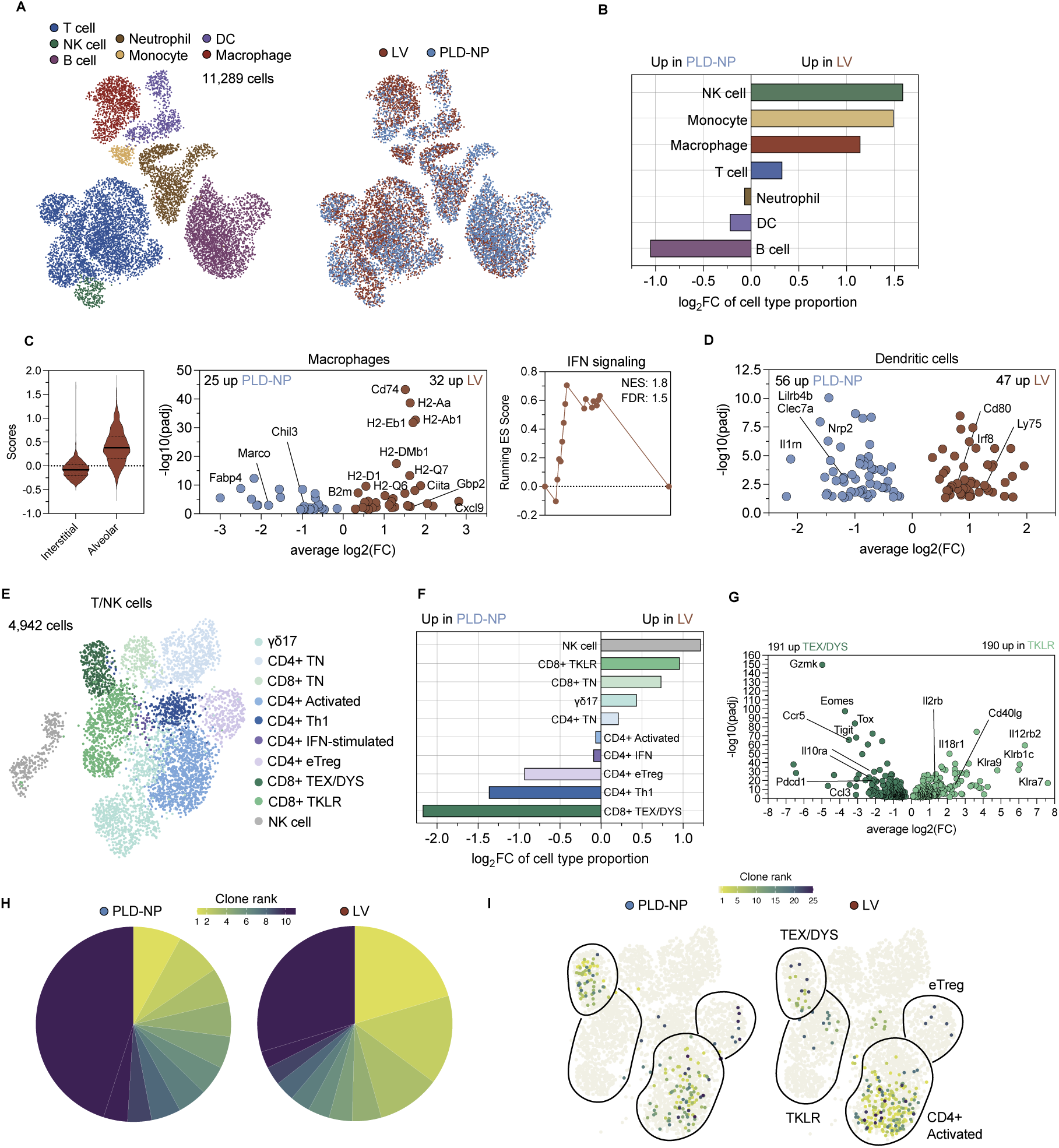
LV instillation drives sustained myeloid remodeling and induces distinct T cell states. (A) UMAP visualization of single-cell transcriptomes of CD45+ cells from LV- and PLD-NP-induced KP tumors labeled by immune cell type (left) and delivery method (right). (B) Abundance of cell populations by delivery method. (C) Left: Interstitial and alveolar scores for the macrophage population. Middle: Differentially expressed genes (DEG) between macrophages from LV-vs. PLD-NP-induced tumors. Right: GSEA enrichment plot for Reactome IFN signaling pathway in LV vs. PLD-NP macrophages. (D) DEG between dendritic cells from LV-vs. PLD-NP-induced tumors. (E) UMAP visualization of single-cell transcriptomes of T and NK cells. (F) Abundance of T and NK cell populations by delivery method. (G) DEG of KLR-like cluster vs. TEX/DYS. (H) Pie charts per delivery method depicting the top 12.5% abundant clones of the TCR repertoire. Wedge sizes reflect each clone’s proportion within the graphed repertoire and are colored by clone rank. (I) Top 25 abundant clones per delivery method projected onto the UMAP space of T cell phenotype depicted in 6E. Clones with rank > 25 are shown in beige.

To assess in vivo biodistribution and persistence, we administered PLD-NPs encapsulating luciferase mRNA I.T. and performed longitudinal IVIS imaging. Luminescent signal remained confined to the lungs and dissipated by 72 hours post-delivery, indicating transient and localized expression (Supp. Fig. 6G). To evaluate tumor initiation efficiency of PLD-NPs, we used KP mice harboring a Cre-inducible TdTomato reporter (KPT), enabling visualization and quantification of tumor cells by microscopy and flow cytometry (Fig. 4A). At 9 weeks post-induction, we observed robust tumor burden in both PLD-NP and PGK-LV-injected mice (Fig. 4B-D, Supp. Fig. 6H). Further, TdTomato^+^ tumors initiated by both PLD-NPs and PGK-LVs expressed NKX2-1, a lineage-defining transcription factor for alveolar type II cells, a major cell of origin of LUAD (Fig. 4E).

The PGK-LV-induced autochthonous model is recognized for mirroring human lung cancer progression and histopathology^14, 17^. Time-course analysis at 4-, 6-, and 9-weeks post-induction (Fig. 4F) revealed that both PGK-LV and PLD-NPs generated tumors with similar histological grades, ranging from grade 1 (atypical hyperplasia) to grade 3 (invasive carcinoma), though PLD-NPs produced fewer lesions overall (Fig. 4G-H). Lesions from PGK-LV-treated mice were more likely to be grade 1 lesions at 9 weeks (Fig. 4H), suggesting the presence of dormant, longer latency lesions, either due to early immune control or due to stable integration and continuous Cre expression in the tested PGK-LV system. Based on these findings we concluded that PLD-NP-induced tumors faithfully recapitulate human LUAD disease progression and histopathology, similar to the well-established PGK-LV system.

To reduce the number of individual lesions and more accurately model human disease, we administered a lower dose of PLD-NPs (2.5 µg → 1.25 µg). Tumor formation was still observed, though with fewer and more focal lesions (predominantly grades 1 and 2) at 9 weeks post initiation (Fig. 4I-J, Supp. Fig. 6I), mimicking the focal nature of human LUAD, which typically presents as discrete lesions. To track tumor kinetics in this lower-dose setting, we performed longitudinal microCT imaging (Fig. 4K). Tumors exhibited slower growth kinetics but progressively expanded over time (Fig. 4K). By 22 weeks post-induction, mice harbored tumors with focal lesions, spanning all histological grades (Fig. 4L-M, Supp. Fig. 6J). Further, 60% of mice had mLN metastases (Supp. Fig. 6K). This low-dose, extended-latency model more closely reflects the clinical course of LUAD, enabling more granular studies of tumor-immune interactions as well as metastatic progression, a hallmark of human LUAD and a major barrier to effective treatment. It is important to note that a similar approach of reducing the amount of LV or adenovirus has been used to generate fewer, focal lesions while allowing for greater metastatic burden^15, 44, 45^.

Finally, we assessed the performance of PLD-NP_Lyoph_ and found that they retained the capacity to initiate tumors, confirming the preserved bioactivity of this platform post long-term storage. However, PLD-NP_Lyoph_ were less efficient at initiating tumors despite similar mRNA encapsulation efficiency (Supp. Fig. 5G), likely requiring higher doses to achieve tumor burdens comparable to freshly prepared PLD-NPs (Supp. Fig. 6L). These data demonstrate that PLD-NP-induced tumors preserved the histologic and molecular characteristics of human LUAD progression, as previously established for PGK-LV models.

### PLD-NPs achieve Cre-mRNA delivery without innate immune transfection and activation

To investigate whether PLD-NPs infect and activate immune cells, we performed immune phenotyping seven days after I.T. administration of PGK-LVs and PLD-NPs in TdTomato reporter mice, where Cre-mediated recombination induces TdTomato expression in transfected cells (Fig. 5A), in the absence of tumors. Since mice cannot form tumors upon Cre delivery, any TdTomato signal detected in immune cells likely reflects true infection rather than uptake of fluorescent tumor cell debris by highly phagocytic APCs such as AMs and DCs.

Both PGK-LV and PLD-NPs transfected comparable proportions of CD45-non-immune cells (Fig. 5B), while PLD-NPs showed significantly reduced infection of CD45+ immune cells in the lung compared to PGK-LVs (Fig. 5C-D). PGK-LVs infected AMs and DCs which was almost entirely abrogated with PLD-NPs (Fig. 5E-F). By minimizing immune cell infection, PLD-NPs help to preserve the fidelity of the LUAD model. Importantly, PLD-NP_Lyoph_ functioned similarly to freshly made PLD-NPs by avoiding AM and DC transfection (Fig. 5B-F).

Given the widespread use of lipid-based nanoparticles (Lipid-NPs) for mRNA delivery, we directly compared Lipid-NP and polymer-based NP systems. Unlayered Lipid-NPs (UL-Lipid-NPs), PLD-layered Lipid-NPs (PLD-Lipid-NPs), and PBAE-based PLD-NPs exhibited comparable size, polydispersity, and in vitro transfection efficiency (Supp. Fig. 7A-C). However, in vivo, both UL-Lipid-NPs and PLD-Lipid-NPs transfected AMs in the lungs, while UL-Lipid-NPs also transfected DCs in the mLN (Supp. Fig. 7D-E). In contrast, PBAE-based PLD-NPs did not transfect these myeloid immune cells (Supp. Fig. 7D-E).

We next assessed DC activation in the lungs and mLN as DCs are crucial for shaping adaptive anti-tumor immune responses. DCs from PGK-LV-treated mice increased in number (Supp. Fig. 8A-B) and displayed robust upregulation of the co-stimulatory markers CD40, CD80, and CD86, as well as the inhibitory ligand PD-L1 (Fig. 5G-I, Supp. Fig. 8C-D), consistent with strong innate immune activation. In contrast, DCs from PLD-NP–treated mice were characterized by significantly lower costimulatory and coinhibitory marker expression (Fig. 5G-I, Supp. Fig. 8A-D).

To test whether the observed differential activation of the myeloid compartment affected tumor dynamics, we infected C57BL/6 mice with PGK-LV-LucOS and subsequently challenged them 1 week later with KP parental cells using tail vein administration. The chosen KP cells lack any known shared antigens including LucOS and Cre (Fig. 5J). In the PGK-LV-LucOS cohort we observed a significant reduction in tumor burden (Fig. 5K), suggesting that PGK-LV administration alters tissue-resident immune cell populations and function in a manner that influences tumor growth mostly in an antigen independent fashion. Consistent with prior data demonstrating minimal myeloid activation by PLD-NP (Fig. 5. E-I), PLD-NP administration did not affect tumor growth in mice challenged with KP lung tumors (Fig. 5L-M). These data demonstrate that while PGK-LVs are powerful tools for tumor initiation in the lung, they can trigger strong activation of myeloid immune cells, impacting tumor growth and progression and might confound the interpretation of tumor-immune interactions. These limitations underscore the benefit of a virus-free platform for tumor induction that maintains genetic precision while avoiding myeloid perturbation.

### Lentiviral Cre delivery and subsequent transformation drive sustained myeloid remodeling and distinct T cell states

To understand how both delivery methods influenced the immune landscape of KP lung tumors, we collected lung tissue 14 weeks after I.T. administration of PGK-LVs and PLD-NPs and performed paired scRNAseq and TCRseq of the CD45+ immune cells from lungs. These experiments were performed in Kras^LSL-G12D/+^;p53^fl/fl^;LSL-SIY mice, in which Cre delivery co-activates expression of the model antigen SIY in transformed cells, with the intent of tracking antigen-specific CD8 T cells. However, SIY-reactive CD8 T cells were undetectable by tetramer staining in both delivery conditions, consistent with prior work showing that thymic expression of SIY induces central tolerance in this system ^46^. Total lung tumor burden was similar in mice treated with either PGK-LVs or PLD-NPs at 13 weeks post initiation (Supp. Fig. 9A). After alignment, quality control, normalization, and dimension reduction (see Methods), we obtained over 11,000 cells with high-quality gene expression profiles. Cells were clustered and labeled with 7 broad immune cell categories, each defined by canonical gene expression (Supp. Fig. 9B, Table 1). Figure 6A illustrates cells labeled by delivery method or immune cell type. We discovered that the proportion of NK cells, monocytes, and macrophages were 2-fold higher in tumors initiated by PGK-LV *vs.* PLD-NP (Fig. 6B, Supp. Fig. 9C). PLD-NP-induced tumors were enriched for B cells; however, differentially expressed gene analysis (DEG) of B cells between the two delivery methods indicate no clear differences in activation or antigen presentation (Supp. Fig. 9D).

Sequenced macrophages scored highly for an alveolar macrophage signature score^47^ (Fig. 6C). Macrophages from PGK-LV-induced tumors had higher expression of genes associated with MHC-I presentation (*B2m, H2-D1, H2-Q6, H2-Q7*), MHC-II presentation (*Ciita, Cd74, H2-Aa, H2-Ab1*, *etc.*), and an IFNγ response (*Gbp2, Cxcl9*). In addition, Gene Set Enrichment Analysis (GSEA) using Reactome pathways revealed an upregulation of IFN signaling in PGK-LV-*vs.* PLD-NP-induced tumors. Lastly, DEG of dendritic cells suggested elevated levels of cross-presentation (*Irf8, Ly75, Cd80*) in the PGK-LV setting versus higher immune regulation in the PLD-NP setting (*Lilrb4b, Il1rn, Nrp2*, *etc.*) (Fig. 6D). These results suggest that compared to PLD-NP, PGK-LV-induced tumors trigger and maintain an anti-viral immune response.

We hypothesized that differences in myeloid magnitude and activation would influence the resulting T cell response. To explore this, T and NK cells were labeled as 10 distinct phenotypes (Fig. 6E, Supp. Fig. 9E-F, Table 2). There were two times higher effector regulatory T cells (eTregs) in PLD-NP *vs.* PGK-LV-induced tumors (Fig. 6F). Interestingly, each delivery method was enriched for a distinct CD8+ effector T cell phenotype, KLR-like in PGK-LV (*Klra7, Klrd1, Klrk1, Il12rb2*) and exhausted/ dysfunctional (TEX/DYS) in PLD-NP (*Gzmk, Tox, Tigit, Entpd1*). DEG between these two populations revealed upregulation of KLR receptors (*Klra7, Klra9, Klrb1c*) and IL-12/IL-18 receptors (*Il12rb2, Il18r1*) in KLR-like T cells and upregulation of canonical exhaustion (*Tox, Tigit, Pdcd1*) and effector markers (*Gzmk, Eomes*) in TEX/DYS (Fig. 6G).

Figure 6H shows the size of the top 10 largest clones for each delivery method, revealing greater clonal expansion among T cells in PGK-LV-induced tumor lesions. Notably, the largest clone in the PGK-LV condition was 2.5 times larger than the largest clone in the PLD-NP condition. Interestingly, across both delivery methods, the top 25 clonotypes were predominantly composed of activated CD4+ T cells (Fig. 6I and Supp. Fig. 9G). Similar to the CD8+ T cell population, CD4+ T cells from PGK-LV-induced tumors showed greater clonal expansion within the Th1-like T cell state (Fig. 6F-I). In contrast, CD4+ T cells from PLD-NP-induced tumors showed greater expansion of TEX/DYS and eTreg states (Fig. 6F-I), further highlighting qualitative differences in the tumor-infiltrating T cell populations generated by the two delivery methods. Notably, nearly half of the top 10 clonotypes in PLD-NP-induced tumors were TEX/DYS (Supp. Fig 9G). Overall, our scRNA-seq/TCR-seq analysis demonstrates that the method used to induce KP tumors substantially influenced the resulting T cell response. PGK-LV-induced tumors were associated with T cell populations dominated by a KLR-like phenotype, whereas PLD-NP-induced tumors exhibited a greater representation of exhausted/dysfunctional T cell states.

## Discussion

GEMMs have been instrumental in elucidating tumor biology and immune responses, yet if the model depends on viral recombinase delivery the immunogenicity associated with viral particles can alter immune response, particularly within myeloid cell populations. While approaches such as tissue-specific promoters, or microRNA-based tissue restriction have been successfully used to mitigate expression of transgenes in non-target cells, innate immune cell sensing of viral particles remains a problem of most viral-based models. Here, we address this critical gap in preclinical modeling of lung cancer by developing a non-viral, NP-based delivery system that enables precise, epithelial-specific oncogene activation and tumor suppressor deletion without confounding innate immune cell activation.

By delivering Cre mRNA via the PLD-NP platform, we initiated lung tumors in KP mice that mirror the histopathological progression, latency, and metastatic behavior of human LUAD. As seen with low viral-titer administration for tumor initiation, low dose PLD-NPs produce tumors with spatially discrete focal lesions, offering a powerful tool to study clonal relationships between individual tumor lesions. PLD-NPs avoid AM and DC transfection and activation, two populations that are readily infected and activated by lentiviral and adenoviral vectors, even when tissue specific promotors are used^10, 18–20^. Bypassing this early inflammatory perturbation alters the tumor immune microenvironment. Although exact dosage comparisons between PGK-LV and PLD-NP are challenging and late-stage tumor burdens were not proactively matched, both methods demonstrated comparable early Cre delivery to lung epithelial cells in tdTomato^LSL^ mice and resulted in similar tumor burdens at most analyzed time points. Nevertheless, it is possible that the observed transcriptional differences in leukocytes from tumors are partly due to disparities in number of lesions, tumor grade, or sustained Cre expression by PGK-LV.

The modularity of our NP design also provides exciting opportunities for further functionalization. Our choice of Poly2, a well-characterized PBAE, is grounded in its high transfection efficiency, biocompatibility, and chemical tunability. PBAE backbones can be readily modified to introduce functional moieties such as targeting ligands, nuclear localization signals (NLS), or even covalently linked antigens^48^. Moreover, LbL assembly permits additional surface modifications that can enhance tissue targeting or immune evasion. While some in vitro studies suggest that Poly2 can stimulate DCs or macrophages^49, 50^, our in vivo findings indicate that LbL coatings like PLD confer stealth properties, minimizing innate immune cell activation and improving the bio interface of polymeric delivery systems.

Despite its advantages, a PLD-NP approach for tumor initiation has limitations. Because PLD-NPs rely on mRNA delivery, they do not permit stable integration of neoantigens into tumors. One potential solution is the use of the NINJA mouse model^23^, which allows for Cre mediated expression of neoantigens in lung cancer models and PLD-NP mediated Cre delivery in this setting would likely minimize innate immune activation while preserving the capacity to track tumor-specific lymphocytes. Moving forward, PLD-NPs can be engineered to deliver transposons, facilitating exogenous neoantigen integration in-situ^48^. Alternatively, this platform could be adapted to deliver pegRNAs to the newly developed prime editor-compatible KP mice for precise modeling of a variety of endogenous antigens^51^. Precedent for this approach comes from recent work demonstrating that cationic PBAE-based nanoparticles can achieve in vivo genome editing in the lung following intratracheal administration^52^. Although that study did not evaluate immune cell infection or activation, it establishes the feasibility of polymer-mediated editing in pulmonary tissues. Furthermore, while our study focused on modeling LUAD, NPs can be engineered to target other tissues (e.g., pancreas, liver) by tuning polyplex surface charge or incorporating tissue-specific targeting ligands, broadening the applicability of this platform beyond lung cancer.

In summary, we describe a virus-free platform for initiating autochthonous LUAD with histopathology, latency, and metastatic behavior comparable to PGK-LV models, but without transduction or activation of myeloid cells within the tumor tissue. The choice of delivery modality has important implications for understanding immune dynamics in tumor outgrowth. Viral delivery for transformation remains a well-validated option which in addition offers flexibility in neoantigen instillation. In contrast, the PLD-NP based approach described here drastically minimizes infection and activation of myeloid cells and makes the tumor-microenvironment a product of the tumor rather than the vehicle used for tumor initiation.

## Materials and Methods

### Mice

Female and male Kras^LSL-G12D/+^ ; Trp53^fl/fl^ (KP) mice on a C57BL/6 background were bred and maintained in-house. B6.Cg-Gt(ROSA)^26Sortm14(CAG-tdTomato)Hze^/J (strain #007914) (T) and B6.129S4-Gt(ROSA)^26Sortm3(CAG-luc)Tyj/J^ (strain #009044) (SIY) mice were purchased from the Jackson Laboratories and bred to KP mice to generate KPT and KPSIY mice. Female C57BL/6 mice (6-14 weeks) were purchased from Jackson Laboratories. Mice were maintained under specific pathogen-free conditions at the Koch Institute animal facility, and all animal procedures were approved by the Committee on Animal Care (CAC/IACUC) at MIT.

### Cell lines

KP, KP.SIY, and the Green-Go reporter cell lines^12^ were cultured in DMEM (Gibco) supplemented with 10% heat-inactivated fetal bovine serum, 10 mM HEPES (Gibco), 1X non-essential amino acids (Gibco), 1% Penicillin/streptomycin (Gibco). Cells were maintained at 37°C, 5% CO₂ and passaged using 0.05% trypsin. Mycoplasma testing was routinely performed.

### Virus production and injection

Cre-expressing lentiviruses were generated by transfecting Lenti-X cells with a transfer plasmid, psPAX2 (gift from Didier Trono—Addgene plasmid, 12260) and pMD2.G (gift from Didier Trono— Addgene plasmid, 12259) at a 4:3:1 ratio using polyethylenimine. Viral supernatants were filtered (0.45 μm), ultracentrifuged at 25,000 rpm for 1.5 h, and resuspended in Opti-MEM. Titers were measured using Green-Go cells and 25,000 transforming units were delivered I.T. in 50 µl of Opti-MEM.

Ad5-SPC-Cre^53^ was purchased from the University of Iowa Viral Vector Core. For in vivo experiments, 2.5 x 10^8^ plaque forming units (p.f.u.) of Ad5-SPC-Cre were delivered I.T. in 50 µl Opti-MEM following calcium chloride precipitation.

### PBAE synthesis

The PBAE polymer Poly2 was synthesized as previously described^33^. Poly2 was stored at room temperature under vacuum in a desiccator. The polymer was resuspended at 2 mg/mL in *d-* chloroform*-d*; proton nuclear magnetic resonance (^1^H NMR) and correlation spectroscopy (COSY) NMR spectra were recorded using a three-channel Bruker Avance Neo spectrometer 500.34 MHz equipped with a 5 mm liquid-nitrogen cooled Prodigy broad band observe cryoprobe. Spectra were processed using MestReNova v14.2 (Mestrelab Research). Peaks were identified based on both 1D and 2D spectra. The number average molecular weight (*M_n_*), the weight average molecular weight (*M_w_*), and the polydispersity index (PDI) of the polymer were measured using the Viscotek GPCmax VE 2001 system equipped with the Ultrahydrogel^TM^ 6 x 40 mm Guard Column and the Ultrahydrogel^TM^ 250 7.8 x 300 mm Column. Sample was detected with the Viscotek VE 3580 Refractive Index Detector. The mobile phase was 1M sodium acetate pH 3 buffer, eluted at 0.6 mL/min. The column temperature was 35°C. A calibration curve was generated using polyethylene glycol standards (Agilent, PL2080-0201).

### PLD-polyplex formation

All solutions were sterile filtered in 0.2 µm cellulose acetate filters before use. Poly2 was dissolved in 25 mM sodium acetate (NaAc) (pH 5.2) by sonicating for 3 minutes then shaking rapidly for 1h. Cre mRNA (5moU, TriLink Biotechnologies) was thawed to room temperature, then dissolved in 25 mM NaAc at 0.05 or 0.1 mg/mL. To form polyplexes, Poly2 solution was rapidly pipetted into an equal volume of mRNA solution in DNA LoBinds, followed by 30 min incubation at room temperature. The weight ratio of Poly2 to mRNA was varied. Poly-L-aspartic acid (PLD) (PLD100, Alamanda Polymers) was dissolved at 20 mg/mL in ultrapure water, by vortexing rapidly and sonicating for 10 min. To layer the polyplexes, 6 weight equivalents of PLD to Poly2 were dissolved further in ultrapure water (upon optimization), or in a solution of 50 mM HEPES, 25 mM NaCl (prior to optimization). In DNA Lo-Binds, polyplexes were added over PLD solution and mixed by rapidly pipetting. PLD-polyplexes were incubated at room temperature for 5 min, shaking. Poly-L-glutamic acid (PLE) (PLE100, Alamanda Polymers), hyaluronic acid (HA) (HA-20k, Lifecore Technologies), and poly(acrylic acid) (PAA) (15kDa, Sigma-Aldrich) were dissolved in ultrapure water and tested for layering at different weight ratios.

### Polyplex concentration

Unlayered polyplexes were concentrated by dialysis against a PEG-20K solution. PEG-20K (Sigma Aldrich 81300) was dissolved in de-ionized water at 0.4 g/L, by gentle heating and physical agitation. Slide-a-Lyzers, 3.5K MWCO (Thermo Scientific 69550) were cleaned with 2 volumes of water, then reservoirs were filled with PEG-20K. Unlayered polyplexes were loaded into Slide-a-Lyzers, and allowed to equilibrate against PEG-20K shaking at 300 rpm, at room temperature. PEG-20K in reservoirs was fully replaced every 1-2 h. Dialysis continued until sample volume reduced by approximately two-fold.

### Lipid nanoparticle preparation

Unlayered and layered lipid nanoparticles (LNPs) were prepared as previously described^54^. LNPs encapsulating Cre mRNA were composed of 50 mol % ALC-0315 (Avanti #890900), 38.5 mol % cholesterol (Avanti #7000000), 1.5 mol % DMG-PEG-2000 (Cayman Chemical Co 33945), and 10 mol % DOPE (Avanti #850725).

To layer LNPs, PLD100 was dissolved at 10 mg/mL in ultrapure water, by vortexing rapidly and sonicating for 10 min. LNPs were layered with 2 weight equivalents of PLD, dissolved into 5 mM HEPES. After both formulation and layering, LNPs were purified and concentrated via three washes in water, in Amicon Ultra-4 100K MWCO ultracentrifugal filter units.

### NP characterization

#### Dynamic light scattering

Hydrodynamic diameters, polydispersity index, and zeta potential of polyplexes were measured in a Malvern Zetasizer Pro (Malvern Panalytical). Unlayered polyplexes were measured in NaAc 25 mM; PLD-polyplexes and concentrated unlayered polyplexes were measured in ultrapure water. Measurements were done at 25°C, with a red laser (λ=633nm) and a detection angle of 173°.

#### Encapsulation efficiency and concentration

mRNA encapsulation was confirmed with fluorescent plate-based Quant-it RiboGreen RNA assays and DNA gels. In a Nunc F96 MicroWell Black polystyrene plate, 5 μL of polyplexes were incubated in 45 μL of either 1× TE or 0.5x SDS in TE. Samples were shaken at 130 rpm, at 37 °C, for 10 min. RiboGreen reagent was diluted 200-fold in 1× TE and protected from light. Samples were then mixed with 50 μL of the diluted RiboGreen reagent. Then, samples were shaken at 300 rpm at room temperature for 5 min, protected from light. Fluorescence intensities were read immediately on a Tecan M1000 plate reader, at an excitation of 485 nm and emission of 525 nm. Encapsulation was calculated as (Fluorescence of SDS polyplexes−Fluorescence of TE polyplexes)/(Fluorescence of SDS polyplexes).

Polyplex samples and free mRNA controls were loaded onto E-Gel EX 1% agarose gels (Thermo Scientific G401001). Gels were run on an Invitrogen E-Gel Power Snap Electrophoresis system and imaged on the ChemiDoc MP imaging system (BioRad 12003154).

#### Cryo transmission-electron microscopy

In sample preparation for cryo-electron transmission microscopy, 3 µL of the sample in buffer solution was applied to copper grids coated with a continuous carbon film. The grids were pre-treated with oxygen plasma using a Solarus 950 Gatan Advanced Plasma System. Excess sample on the grid was gently blotted using the Gatan Cryo Plunge III, followed by rapid plunging into liquid ethane to vitrify the sample. The grid was then mounted on a Gatan 626 single tilt cryo-holder, which was subsequently inserted into the TEM column. Both the specimen and the holder tip were maintained at cryogenic temperatures with liquid nitrogen to ensure preservation throughout the transfer and imaging process. Imaging was performed on a JEOL 2100 FEG microscope using a minimum dose method to reduce electron beam damage to the sample. The microscope was operated at 200 kV, with magnifications between 10,000x and 60,000x to evaluate particle size and distribution. All images were captured using a Gatan 2k x 2k UltraScan CCD camera.

### NP lyophilization

Sucrose or trehalose were dissolved in water at 0.5 g/mL with rapid vortexing. NPs at 25 ng/μL mRNA were suspended in 5% sucrose or 10% trehalose via rapid pipetting. Samples were shaken at 300 rpm, room temperature, for 5 min, before they were frozen at -80°C for 2-6 h. Samples were then lyophilized in a Labconco lyophilizer overnight. Lyophilized samples were sealed and stored with desiccant, either at 4°C or -20°C. Lyophilized samples were resuspended at 25 ng/μL in ultrapure water, immediately vortexing for several seconds upon resuspension and characterized via dynamic light scattering.

### Green-Go reporter assay

Green-Go reporter cells were plated at 10,000 cells/well in a clear flat-bottom 96-well plate. 24h after seeding, cells were treated with polyplexes containing Cre mRNA at 100 ng mRNA/well. 72h after dosing, Cre-mediated GFP expression was evaluated via fluorescence microscopy and flow cytometry. Cells were dissociated with 0.05% Trypsin and prepped for flow cytometric analyses as described below. Images of live cells were acquired using the EVOS Cell Imaging System (ThermoFisher Scientific) with a 10X objective and light cubes for GFP (470/525 nm) and brightfield. All images were processed with the FIJI software package^55^.

### Tumor induction

Lung tumors were induced in the KP and KPT mice by I.T. injection of lentivirus expressing Cre (25,000 TU/mouse) or PLD-NPs containing Cre mRNA to anesthetized mice as previously described^17^. For PLD-NP, 2.5 µg of mRNA was injected per mouse (50 µL of NPs at 50 ng/µL). For the PLD-NP low dose and lyophilized setting,1.25 µg of mRNA was injected per mouse (50 µL of NPs at 25 ng/µL). Transplantable lung and flank tumors were generated by injecting 250,000 cells (KP.SIY or KP) via tail vein or subcutaneous injection respectively.

### Tumor burden analysis

Flank tumor size was measured by calculating the area of the subcutaneous tumor (length x width). Hematoxylin and Eosin (H&E) staining of lung tissue sections was used to determine lung tumor burden, and tumor grade using the Aiforia AI-assisted pathology web tool (www.aiforia.com) with the nsclc_v37_noheatmap_20-02-05 algorithm as previously described^45,51^.

### Tissue processing for flow cytometry

To differentiate immune cells in circulation from the ones in the lung tumor parenchyma, we retro-orbitally injected mice with 2 µg of a fluorescence-labeled anti-CD45 antibody diluted in 100 µL of PBS, 3-5 minutes prior to euthanasia. Animals were euthanized via a retro-orbital injection of 100 μL of Ketamine and Xylazine solution (9 mg/ml Ketamine and 1 mg/ml Xylazine diluted in PBS). Lungs, mediastinal lymph nodes, and spleen were collected at different time points and processed according to the following dissociation methods.

#### Lung

To evaluate tumor, epithelial, and endothelial cells in the lung, tissue was minced and enzymatically dissociated by incubating in a solution containing DMEM/F12, HEPES, DNase, collagenase from Clostridium histolyticum (167 μg/mL), and CaCl_2_ (0.06 mM) for 30 minutes at 37°C (600 rpm). Dissociated tissues were passed through a 100 μm strainer and washed with PBS. Cells were treated with ACK buffer for 3 minutes at room temperature to lyse red blood cells, followed by a PBS wash. Tumor cells were resuspended in FACS buffer (PBS, 2% FBS and 2 mM EDTA) and kept at 4°C for downstream analysis.

Lung tissue used for analysis of immune cells was mechanically processed using a gentleMACS octodissociator and enzymatically digested in HBSS (+Ca +Mg) containing 1mg/ml Collagenase D and DNase for 30 minutes at 37C. Dissociated cells were filtered through a 70 μm strainer and washed with RPMI. Whole tissue homogenate was treated with ACK buffer for red blood cell lysis for 3 minutes on ice. Samples were washed with chilled FACS buffer and stored at 4°C for further analysis. For T cell analysis at later time points (9 weeks post tumor induction), dissociated lung samples were resuspended in 10 ml of PBS and layered over 5 ml of Ficoll (GE) and centrifuged at 450 rcf for 30 minutes with zero acceleration and brake. Immune cells located at the interface between PBS and Ficoll were collected and washed with PBS. Dissociated were washed with FACS buffer and stored at 4°C for further analysis.

#### Spleen

Spleens were directly processed by mashing through a 70 μm filter into RPMI, followed by a PBS wash. Spleens were treated with ACK buffer for red blood cell lysis for 3 minutes on ice, then washed with FACS buffer and stored at 4°C for further analysis.

#### Lymph node

Lymph nodes were mechanically processed by smashing them on a scored plate and enzymatically digested in 1mg/ml of Collagenase D in HBSS with calcium and magnesium and 0.5 mg/ml of DNAse for 15 minutes at 37°C. Dissociated cells were passed through a 70 μm filter into RPMI, followed by a PBS and FACS buffer wash and stored at 4°C for further analysis.

### Flow cytometry analysis

Cells were stained in FACS buffer containing a Fixable Viability dye and with anti-CD16/anti-CD32 antibodies to avoid non-specific binding for 15 min at 4°C. Cells were washed twice with chilled FACS buffer and then stained with a mix of fluorescence conjugated antibodies against surface proteins for 30 min at 4°C. Samples were washed with chilled FACS buffers 3 times and then fixed with 0.5% PFA in FACS buffer for 30 min at 4°C. Cells were washed 3 times and when intracellular staining was needed samples were incubated with a mix of fluorescence conjugated antibodies (Supp, Table 1) diluted in FACS buffer overnight at 4°C. Finally, samples were washed 3 times and resuspended in FACS buffer containing Precision Count Beads (BioLegend) to determine total cell count numbers and kept at 4°C until flow cytometry analysis. Flow analysis was performed using a LSR Fortessa cytometer (BD) or a FACS Symphony A3 (BD). Data collected was analyzed using FlowJo v10.5.3 software (TreeStar).

### Tissue processing for histology and immunofluorescence

For paraffin embedded samples, mouse lung and flank tissues were collected at different time points. Lungs were inflated via trachea injection with 10% formalin solution (Sigma). Tissue was fixed in 10% formalin for 72 hours at room temperature. Tissue was incubated in 70% ethanol followed by 95% ethanol, 100% ethanol, and Xylene. Samples were embedded in paraffin and sliced at 5 μm.

For frozen tissue samples, mice were perfused with 10 ml PBS followed by 10 ml of 4% PFA in PBS into the heart’s right ventricle. Lungs were dissected and inflated with 4% PFA via intratracheal injection and incubated in the same fixative for 24 hours. Tissue samples were then incubated for 24 hours in 15% sucrose solution followed by 30% sucrose solution in PBS at 4C. Lung tissue was inflated with 50% optimum cutting temperature compound (OCT) and 50% PBS solution and mounted in 100% OCT and snap-frozen in 2-Methylbutane on dry ice and stored at -80 until sectioning. Frozen samples were sectioned at 10 μm using the Cryostar NX70 (Thermo Scientific). Post-fixation was performed by incubation of tissue sections in ice-cold acetone for 10 min at -20C. Samples were dried for 1 hour and stored at -20°C.

### Histology and immunofluorescence

Paraffin embedded sections were incubated for 20 minutes at 55°C in a slide warmer. Samples were deparaffined by incubation for 3 minutes in Xylene, 3 times, then 3 minutes in 100% ethanol, 1 minute in 95% ethanol, 1 minute 70% ethanol, and 1 minute in running water, twice. For H&E staining, tissue sections underwent the following treatment: immersion in Harris Acidified Hematoxylin for 3 minutes, rinsing in running water for 1 minute, incubation in acid alcohol for 5 seconds, followed by another 1-minute rinse in running water. Next, sections were treated with a bluing solution for 1 minute, rinsed again in running water for 1 minute, and immersed in 95% ethanol for 1 minute. This was followed by a 10-second exposure to Eosin, another 1-minute immersion in 95% ethanol, two 1-minute immersions in 100% ethanol, and three 2-minute immersions in Xylene. Finally, the sections were mounted using Tissue-Tek® Glas™ Mounting Medium (Sakura).

For immunofluorescence staining of paraffin embedded tissue, antigen retrieval was performed by incubating sections in citrate buffer using the 2100 Retriever, following the manufacturer’s instructions (Electron Microscopy Science). Samples were blocked for 2 hours at room temperature with a blocking solution containing Blocker Casein in tris-buffered saline (ThermoScientific, Cat# 37532), 0.3% Triton X-100, and 10% normal donkey or goat serum (Jackson Immunoresearch). Primary antibodies, diluted in the blocking solution, were applied, and the sections were incubated overnight at 4°C in a humidified chamber. After washing with PBS containing 0.1% Triton X-100, samples were incubated for 1 hour at room temperature with secondary conjugated antibodies diluted in the blocking solution. Sections were then washed three times with PBS and treated with TrueBlack for 30 seconds, according to the manufacturer’s protocol (Biotium, Cat# 23007). After additional PBS washes, nuclei were counterstained with DAPI (Sigma). Tissue sections were mounted using ProLong Diamond Antifade Mountant (Thermo Fisher Scientific, Cat# P36970) and allowed to cure for 24 hours before imaging.

### Image acquisition and analysis

Hematoxylin and eosin-stained samples were imaged using the Leica Aperio Slide Scanner (Leica Biosystems) at 20x and visualized using the QuPath software for digital pathology image analysis^56^. Immunofluorescence tissue-stained samples were imaged using the TissueFAXS Plus automated slide scanning system (TissueGnostics GmbH, Austria), which integrates a Zeiss Axio Imager 2 upright microscope with a Märzhäuser motorized stage (Märzhäuser Wetzlar). A Zeiss 20x Plan-Neofluor 0.5 NA air objective was used for image acquisition, with filter sets AF488 (470/24), ET470/30x T495lpxr ET515/30m, AF750 (740/20), ET740/40x T770lpxr ET780lp, and a multiband dichroic qTexasRed/qCy5 (550/15, 640/30) from Chroma Technology, USA. Excitation was provided by a Lumencor Spectra 3 LED light engine (500 mW per channel), and fluorescence images were captured with a Hamamatsu Orca Flash 4.0 V2 cooled digital CMOS camera (C11440-22CU). Images were acquired as z-stacks using a technique called extended focus, with a 3 μm step size that included one step above and one below the focal plane. This method combines all in-focus regions from the z-stack into a single composite image. Image processing, including stitching, was conducted using the TissueFAXS capture/control software.

Density of positive cells was analyzed using TissueQuest image analysis software (TissueGnostics USA) as follows. Lung tumor lesions were designated as Regions of Interest (ROI). Automatic segmentation of nuclei was carried out in TissueQuest using DAPI staining as a marker. The density of positive cells within each ROI was determined by dividing the number of positive events by the corresponding ROI area. Image postprocessing was performed with the image processing package FIJI^55^.

### Rodent µCT

Images were acquired as previously described^51^. Briefly, anesthesia was induced with isoflurane (3%, then maintained at 2.0–2.5% in oxygen—VetEquip) and mice were scanned in a prone position using a Skyscan 1276 (Bruker). The following parameters were used for scanning: 100 kVp source voltage, 200 μA current, 0.5 mm aluminum X-ray filter, 108 ms exposure time and 0.65-degree rotational step size over 360 degrees in a continuous rotation. With 4 × 4 detector binning, the nominal pixel size after reconstruction (Bruker NRecon software) was 40.16 microns. Data were visualized and quantified using ImageJ^57^.

### Paired scRNAseq and TCRseq of CD45+ compartment from KP tumors

#### Sample preparation, library generation, and sequencing

Intravenous leukocytes were labeled with 2μg of anti-CD45 antibody and KP;lox-Stop-lox-SIY mice (n=2 for each condition) were taken down 14 weeks post tumor induction with nanoparticles or lentivirus. Lungs were digested as previously described (GentleMacs with Collagenase D) and CD45 microbeads (Miltenyi) were used to isolate leukocytes from lung tumors. Live, IV-, CD45+ cells were further purified using a FACS ARIA. Library generation and sequencing were performed at the Whitehead Institute Genome Technology Core using on chip multiplexing (10x, 5’ GEX) and a NovaSeq.

#### Data alignment, processing, analysis and availability

Reads were aligned using the nf-core/scrnaseq pipeline v4.1.0 with aligner Cell Ranger multi ^58^. 10X Genomics-provided GRCm39 reference for gene expression and GRCm38 reference for V(D)J alignment were used. Sequencing and alignment quality was evaluated based on the rank plot, mean reads per cell, mapping percentage (gene expression), and percentage of cells with productive V-J spanning pair.

For scRNAseq, low-quality cells were filtered based on number of UMIs, number of genes, and percent mitochondrial UMIs. Doublets were removed using DoubletFinder with an assumed doublet rate of 3% ^59^. Data were normalized using SCTransform, regressing out percent mitochondrial and ribosomal UMIs ^60^. Clustering was performed using Leiden. Differential gene expression was performed using MAST on log-transformed RNA, regressing out percent mitochondrial/ribosomal UMIs and number of UMIs ^61^. Gene set expression analysis was performed using the fgsea package and the Reactome gene sets ^62^. Gene signatures were calculated using AddModuleScore in Seurat.

For TCRseq, contigs were obtained from Cell Ranger multi’s filtered_contig_annotations.csv, which contains only productive calls. Alpha chains were dropped. Beta chains with ≤2 UMIs were also dropped. Cells with 0 or >1 beta chain were filtered. Clonotypes were defined as having identical V gene, J gene, and CDR3 nucleotide sequence. Flexibility was allowed for ambiguous V gene calls. For paired scRNAseq/TCRseq analysis, clonotyping was merged with gene expression data based on matching cell barcode. After quality control and clonotyping we recovered over 4,100 T cells with productive TCRs, 75% of which had paired expression data, resulting in over 3,000 unique clones.

Data are available upon request and will be made publicly available by depositing to GEO.

### Statistical analysis

Statistical analyses were performed using GraphPad Prism (GraphPad). All data are shown as means ± SEM/SD. For flow cytometry, immunohistochemistry, immunofluorescence, and tumor outgrowth studies, statistical analyses were performed with Mann-Whitney U (MWU) test for comparisons of two groups, one-way analysis of variance (ANOVA) for comparisons between multiple groups, or two-way ANOVA for multiple comparisons over time (unless explicitly stated otherwise), with *P < 0.05; **P < 0.01; ***P < 0.001; and ****P < 0.0001. For one-way and two-way ANOVAs, Šídák’s or Tukey’s multiple comparisons tests were utilized as post-tests. For each experiment n numbers are indicated in figure captions.

## Acknowledgements

We would like to thank members of the Spranger, Jacks and Hammond lab for their valuable discussions and contributions. T.A.H. received support from the Ludwig Cancer Center at MIT. N.M.A received fellowship support from F30CA278495 (Ruth Kirschstein National Service Research Award). Z.J.R was supported by American Cancer Society postdoctoral fellowship (PF-24-1244739-01-IBCD). We thank the Koch Institute’s Robert A. Swanson (1969) Biotechnology Center, specifically the Flow Cytometry Core Facility, the Hope Babette Tang (1983) Histology Core Facility, and the Whitehead Institute Genome Technology Core for providing core services. This work was supported by the Koch Institute Support Grant P30-CA014051 from the National Cancer Institute R37CA273819 and the American Cancer Society ACS RSG-23-863228-01-IBCD.

## Author Contributions

V.B., E.T.-M., T.A.H., and S.S. conceived the study and designed the experiments. N.N., C.V., A.B., and T.G.D. designed and synthesized the nanoparticle formulations and performed characterization, lyophilization optimization, and in vitro transfection assays. T.A.H. performed the single-cell RNA-sequencing and TCR-sequencing experiments and analyzed the resulting data, with Z.J.R.. N.M.-A. performed and analyzed µCT imaging. N.M.-A and S.-L.S. provided viral reagents and assisted with experiments. T.J., P.T.H., and S.S. supervised the work and acquired funding. V.B., E.T.-M., T.A.H., and S.S. wrote the manuscript with input from all authors. All authors read and approved the final manuscript.

