## Supplementary material for "Modeling lung adenocarcinoma using layer-by-layer nanoparticles mitigates innate immune cell activation": Supp fig.

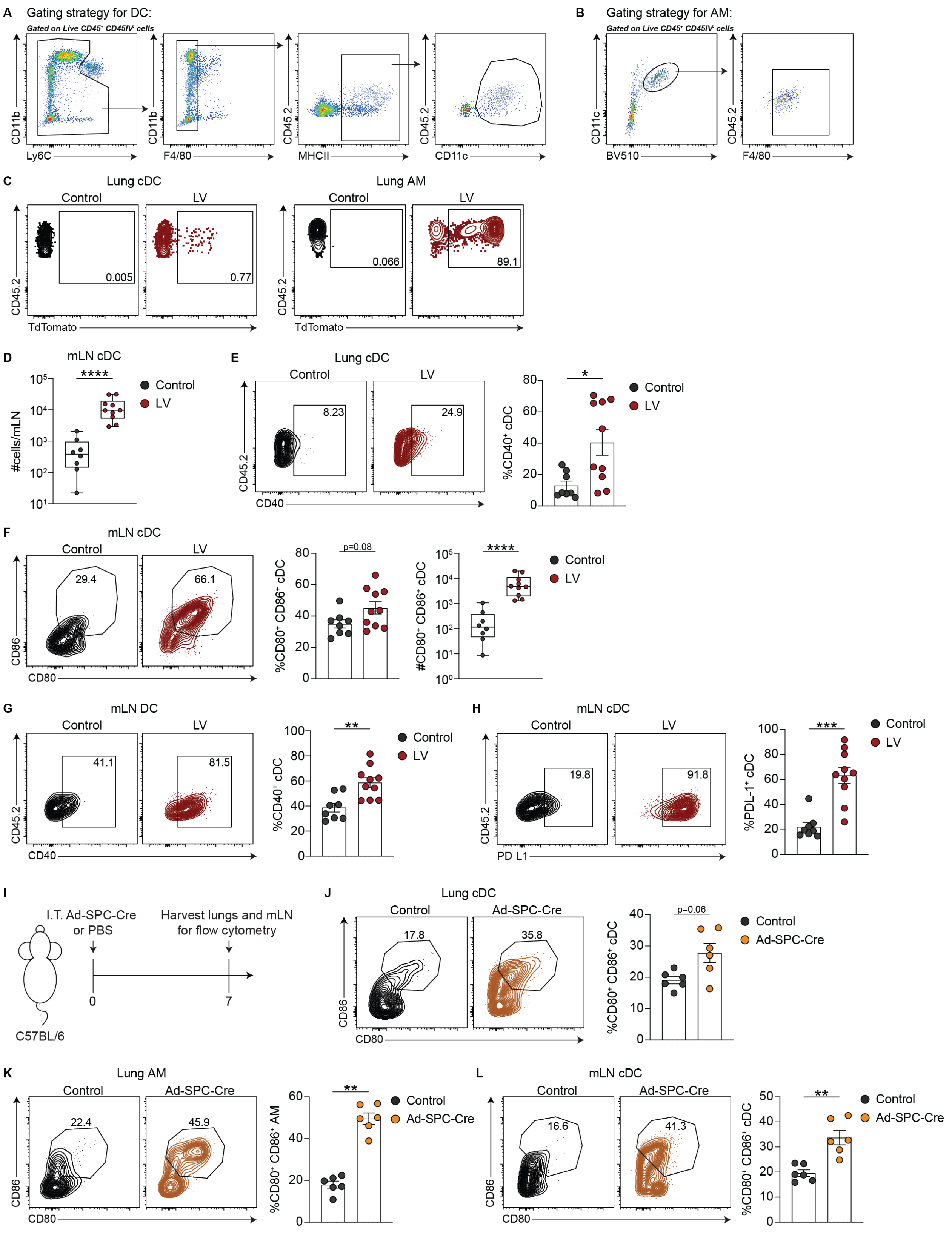


**Supplemental Figure 1.** A-B) Flow cytometric gating for dendritic cells (A) and alveolar macrophages (B). C) Representative flow plots of TdTomato expression by lung DC (left) and

AM (right). D) Number of DCs in the mLN, control n=8, LV n=10. E) Flow plots (left) and quantification (right) of CD40 expression on lung DCs, control n=8, LV n=10. F-H) Flow plots (left) and quantification (right) of CD80,CD86 expression, frequency and numbers (F), CD40 expression (G), and PD-L1 expression (H) on mLN DCs, control n=8, LV n=10. I) Experimental schematic showing I.T. administration of adenovirus encoding SPC-Cre (Ad5-SPC-Cre) to C57BL/6 mice. J-L) Representative flow plots (left) and quantification (right) of CD80, CD86 expression on lung cDC (J) lung AM (K), and mLN cDC (L), n=6. P-values calculated with Mann-Whitney U test. Data shown as mean ± SEM for all quantification.

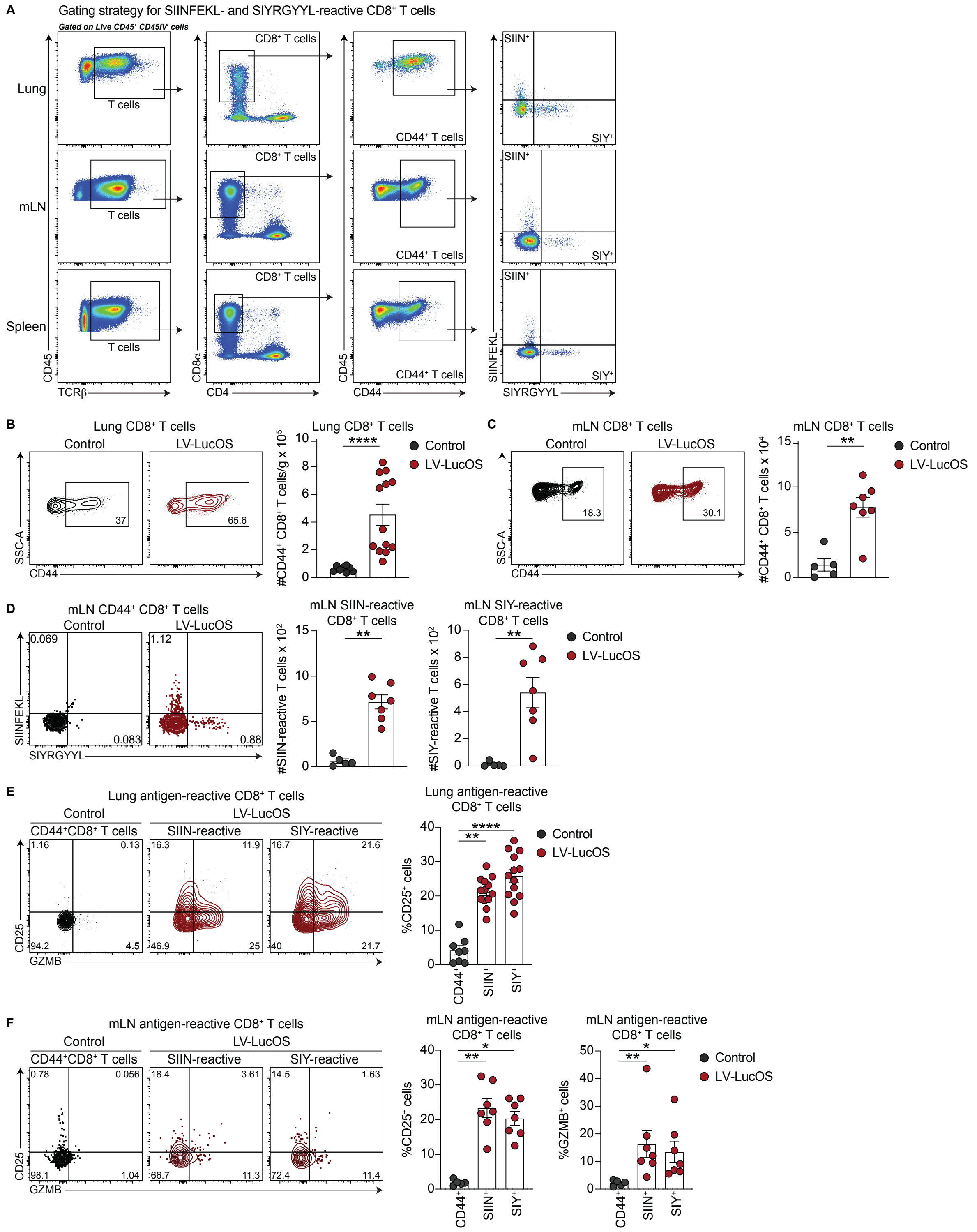


**Supplemental Figure 2.** A) Flow cytometric gating strategies for SIINFEKL- and SIYRGYYL-reactive $CD8^+$ T cells in the lung (top), mLN (middle), and spleen (bottom). B) Representative flow plots (left) and quantification (right) of CD44 expression by $CD8^+$ T cells in the lungs, control n=8, LucOS n=13. C) Representative flow plots (left) and quantification (right) of CD44 expression by $CD8^+$ T cells in the mLN, control n=5, LucOS n=7. D) Representative flow plots (left) and quantification (right) of number of SIIN- and SIY-reactive $CD8^+$ T cells in the mLN of LucOS-infected mice, control n=5, LucOS n=7. E) Representative flow plots of CD25 and GZMB expression by SIIN- and SIY-reactive $CD8^+$ T cells in the lung (left) and quantification of CD25 expression by these cells (right), control n=8, LucOS n=13. F) Representative flow plots of CD25 and GZMB expression by SIIN- and SIY-reactive $CD8^+$ T cells in the mLN (left) and quantification of CD25 and GZMB expression by these cells (right), control n=5, LucOS n=7. For B-D, P-values calculated with Mann-Whitney U test. For E-F, P-values calculated with one-way ANOVA. Data shown as mean ±SEM.

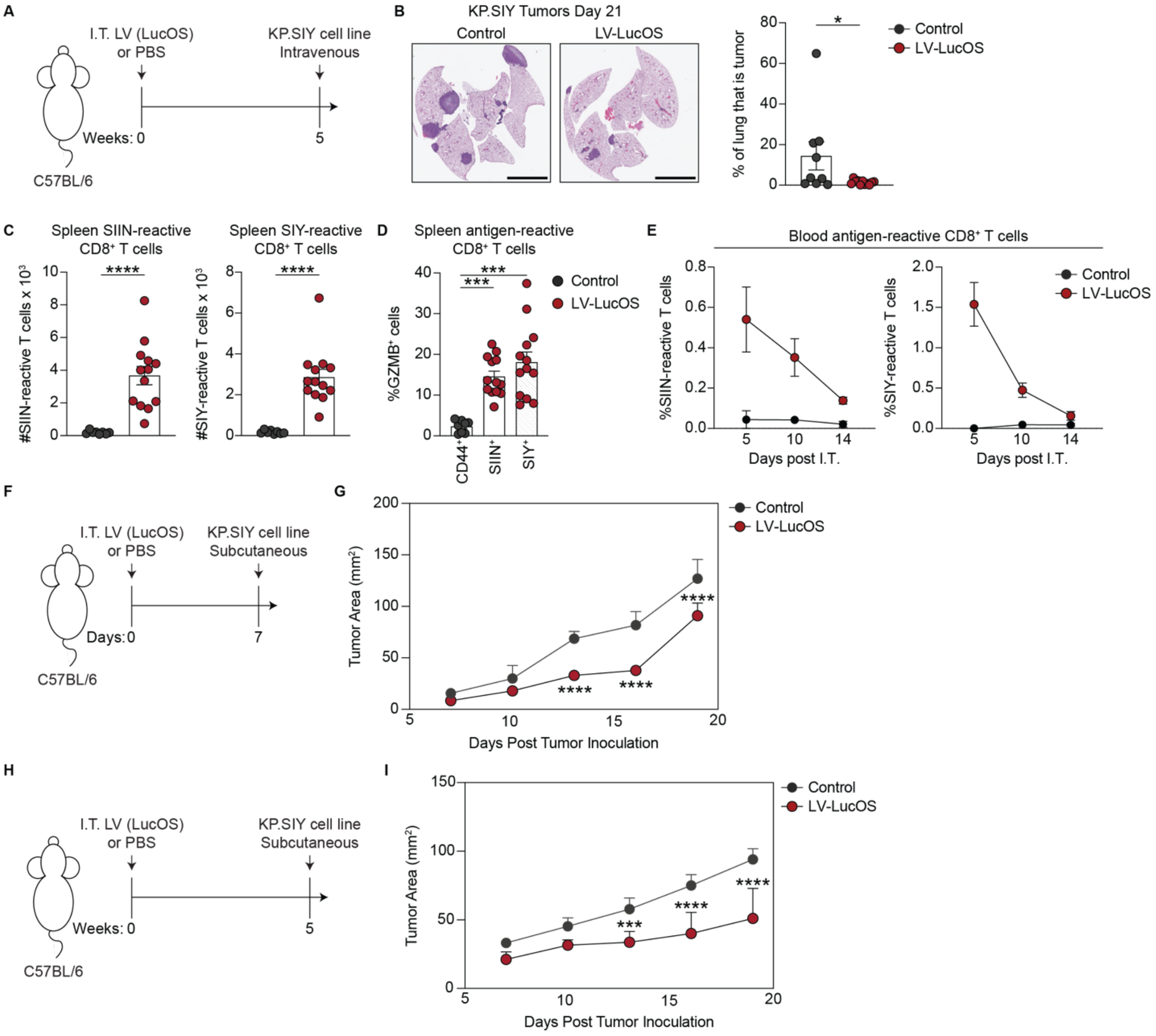


**Supplemental Figure 3.** A) Experimental scheme showing I.T. LucOS delivery followed by I.V. challenge with KP.SIY tumor cells in the memory context. B) Representative H&E staining (left) and quantification (right) of lung tumor burden 21 days post KP.SIY injection, control n=9, LucOS n=8. Scale bar = 4000μM. C) Number of SIINFEKL- and SIYRGYYL-reactive CD8$^+$ T cells in the spleen, control n=8, LucOS n=13. D) Percent of GZMB$^+$ SIIN- or SIY- reactive CD8$^+$ T cells in the spleen, control n=8, LucOS n=13. E) Percent of SIINFEKL- and SIYRGYYL-reactive CD8$^+$ T cells in the blood, control n=6, LucOS n=6. F) Experimental scheme showing I.T. LucOS delivery followed S.Q. challenge with KP.SIY tumor cells in the acute setting. G) KP.SIY flank tumor outgrowth, control n=8, LucOS n=8. H) Experimental scheme showing I.T. LucOS delivery followed S.Q. challenge with KP.SIY tumor cells in the memory setting. I) KP.SIY flank tumor outgrowth, control n=8, LucOS n=8. For B-C P-values calculated with Mann-Whitney U test. For D, , P-values calculated with one-way ANOVA. For G,I, P-values calculated with two-way ANOVA.

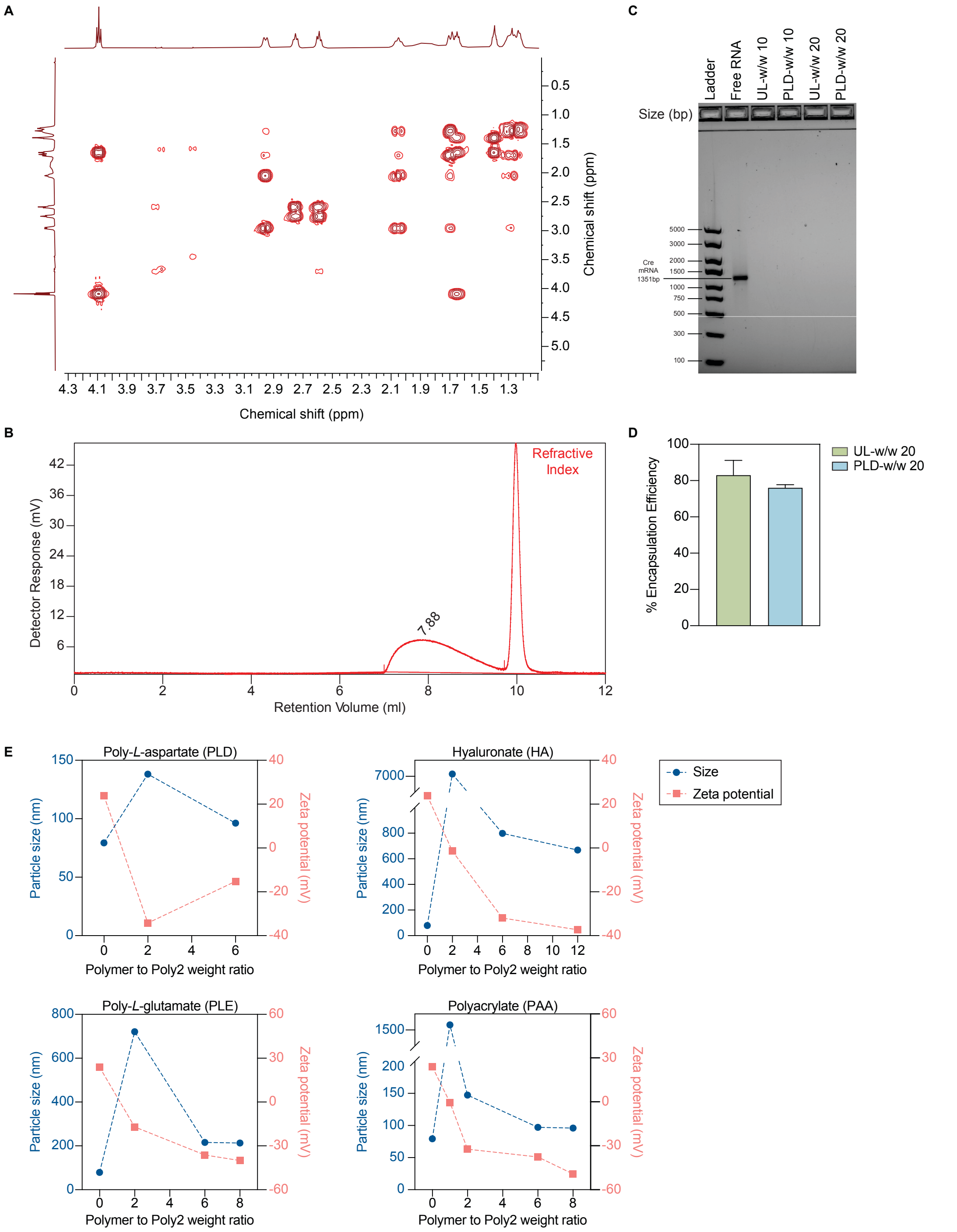


**Supplemental Figure 4.** A) 1H COSY NMR spectrum of Poly-2 in deuterated chloroform ($CDCl_3$). B) Gel permeation chromatography trace of Poly-2. C) 1% agarose gel of polyplexes, in comparison to DNA ladder and free mRNA. D) mRNA encapsulation efficiency using RiboGreen Assay of UL- and PLD-NPs formulated at Poly-2:mRNA weight ratios of 20. N=2–4 replicates. E) Hydrodynamic diameters and zeta potential of Poly-2 polyplexes, layered with varying weight ratios of poly-L-aspartic acid, hyaluronic acid, poly-L-glutamic acid, and poly(acrylic acid). n=1 batch replicate, 3 technical replicates each. Data shown as mean.

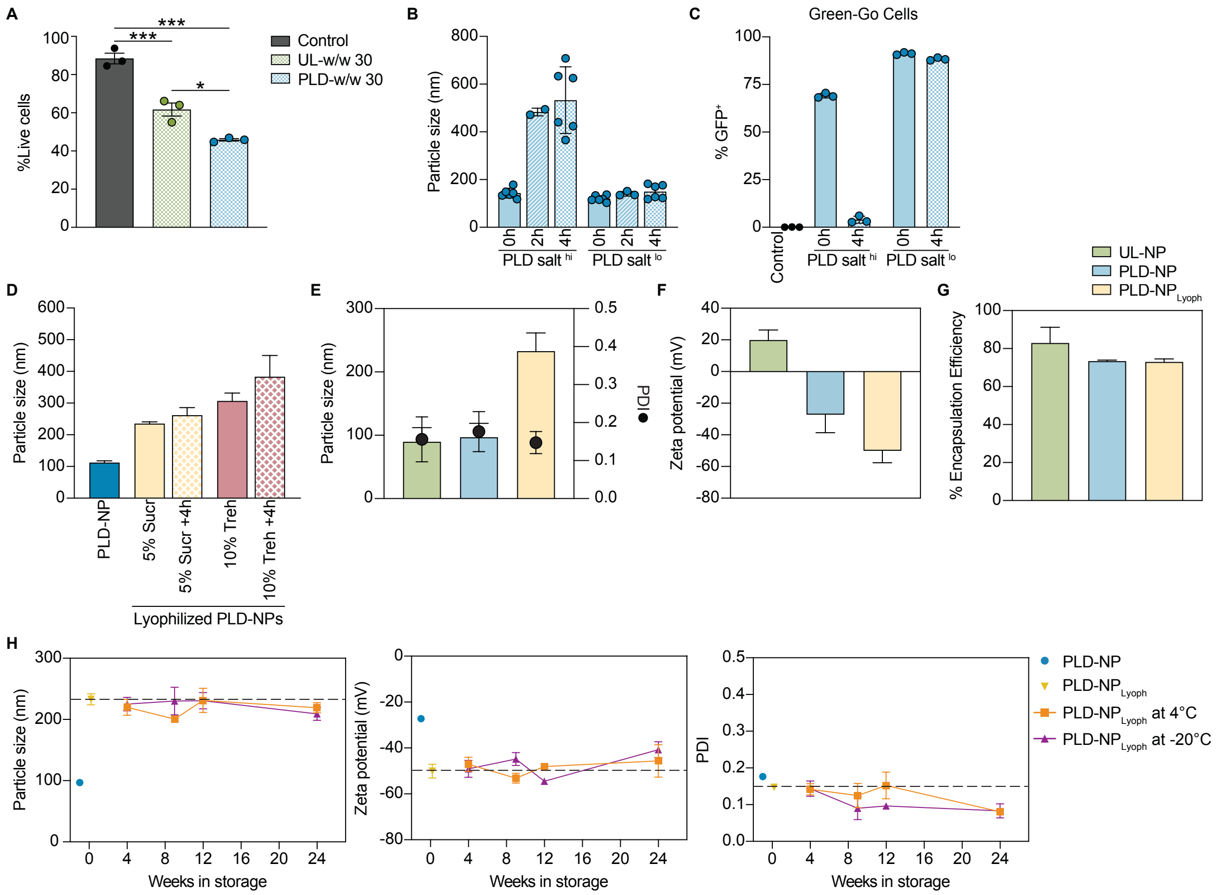


**Supplemental Figure 5.** A) Percentage live cells upon treatment with UL-NPs or PLD-NPs at w/w 30, in comparison to untreated cells. N=1 biological replicate, 3 technical replicates each. B) Hydrodynamic diameter of PLD-NPs layered in either a solution of 50 mM HEPES, 25 mM NaCl (PLD salt $^{hi}$) or ultrapure water (PLD salt $^{lo}$), over a period of 4 hours. N=1-2 batch replicates, 2-3 technical replicates each. C) Percentage GFP-positive Green-Go cells upon treatment with PLD-NPs layered in either a solution of 50 mM HEPES, 25 mM NaCl (PLD salt $^{hi}$) or ultrapure water (PLD salt $^{lo}$). NPs were dosed either 0h or 4h after formulation. N=1 biological replicate, 3 technical replicates each. D) Hydrodynamic diameter of PLD-NPs, either freshly prepared, or lyophilized in each of 5% sucrose (Sucr) or 10% trehalose (Treh), 0 and 4h after resuspension. N=2-3 batch replicates with 3 technical replicates each. E) Hydrodynamic diameter and F) zeta potential of UL-NPs, freshly prepared PLD-NPs, and lyophilized and resuspended PLD-NPs. For UL-NPs, n=76 batch replicates with 1-3 technical replicates each; for freshly prepared PLD-NPs, n= 64 batch replicates with 1-3 technical replicates each; for lyophilized and resuspended PLD-NPs, n=10 batch replicates with 3 technical replicates each. G) mRNA encapsulation efficiency of UL-NPs, freshly prepared PLD-NPs, lyophilized and resuspended PLD-NPs, and stored at 4°C or -20°C for 4 weeks. N=2–4 replicates. H) Hydrodynamic diameter, zeta potential, and polydispersity index (PDI) of freshly prepared PLD-NPs, or lyophilized PLD-NPs stored at either 4°C or -20°C for up to 24 weeks. N=1-3 batch replicates, 3 technical replicates each. All data shown as mean ± SD.

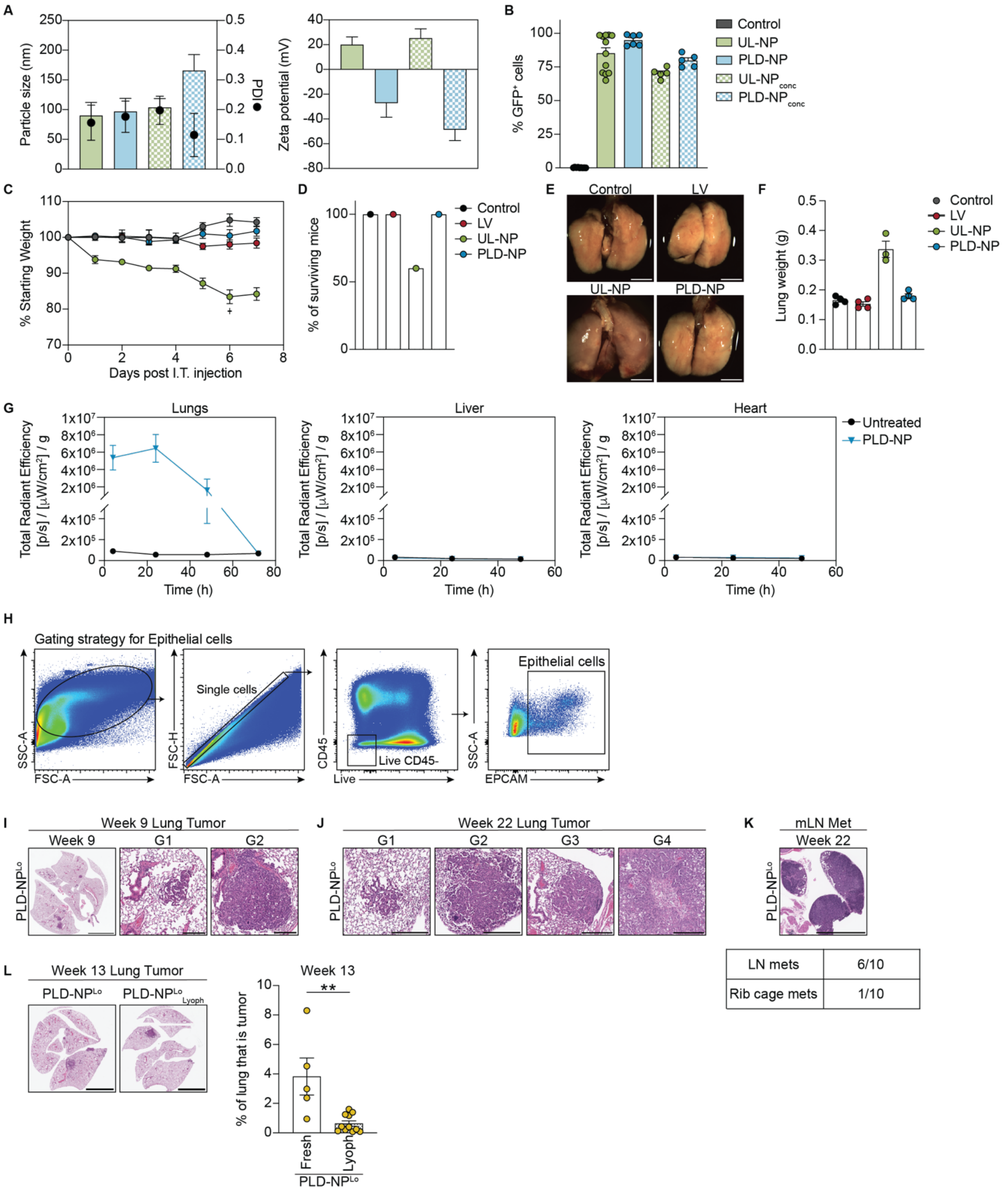


**Supplemental Figure 6.** A) Hydrodynamic diameter, PDI, and zeta potential of concentrated UL- and PLD-NPs. B) Percent of GFP$^+$ Green-Go cells 72hr after transfection. For control, n=5, UL-NP n=12, PLD-NP n=6, UL-NP$_{conc}$ n=5, PLD-NP$_{conc}$ n=5. C) Percent starting weight of mice injected I.T. with PBS, LV, UL-NP, or PLD-NP. For control n=4, LV n=4, UL-NP n=5, PLD-NP n=4. $^+$ indicates the death of 2 mice due to toxicity. E) Representative bright-field and fluorescent images of whole lungs 7 days post I.T. Scale bar = 500 pixels. F) Lung weight 7 days post I.T. For control n=4, LV n=4, UL-NP n=3, PLD-NP n=4. G) Quantification of total radiant efficiency at multiple time-points post I.T. injection to C57BL/6 mice of PLD-NPs encapsulating luciferase mRNA in the lungs (left), liver (middle), and heart (right), n=2-4 biological replicates for untreated mice and n=4-6 biological replicates for PLD-NPs, per time-point. H) Flow cytometric gating strategy for epithelial cells in the lung. I) Representative H&E-stained lung sections, showing tumor burden and histologic grade at 9 weeks in mice with PLD-NP$^{lo}$ initiated tumors. Scale bar for whole lung = 4000μM; scale bar for grades = 300μM. J) Representative H&E-stained lung sections across, showing histologic grade at 22 weeks in mice with PLD-NP$^{lo}$ initiated tumors. Scale bar = 300μM. K) Representative H&E-stained mLN section (top) and quantification of proportion of mice with metastases (bottom) at 22 weeks post tumor initiation with PLD-NP$^{lo}$. Scale bar = 1000μM. L) Representative H&E-stained lung sections (left) and quantification (right) of tumor burden at 13 weeks post tumor initiation with freshly prepared and lyophilized PLD-NPs, fresh n=5, lyoph n=11. Scale bar = 4000μM. P-values calculated using Mann-Whitney U test. All data shown as mean ± SEM.

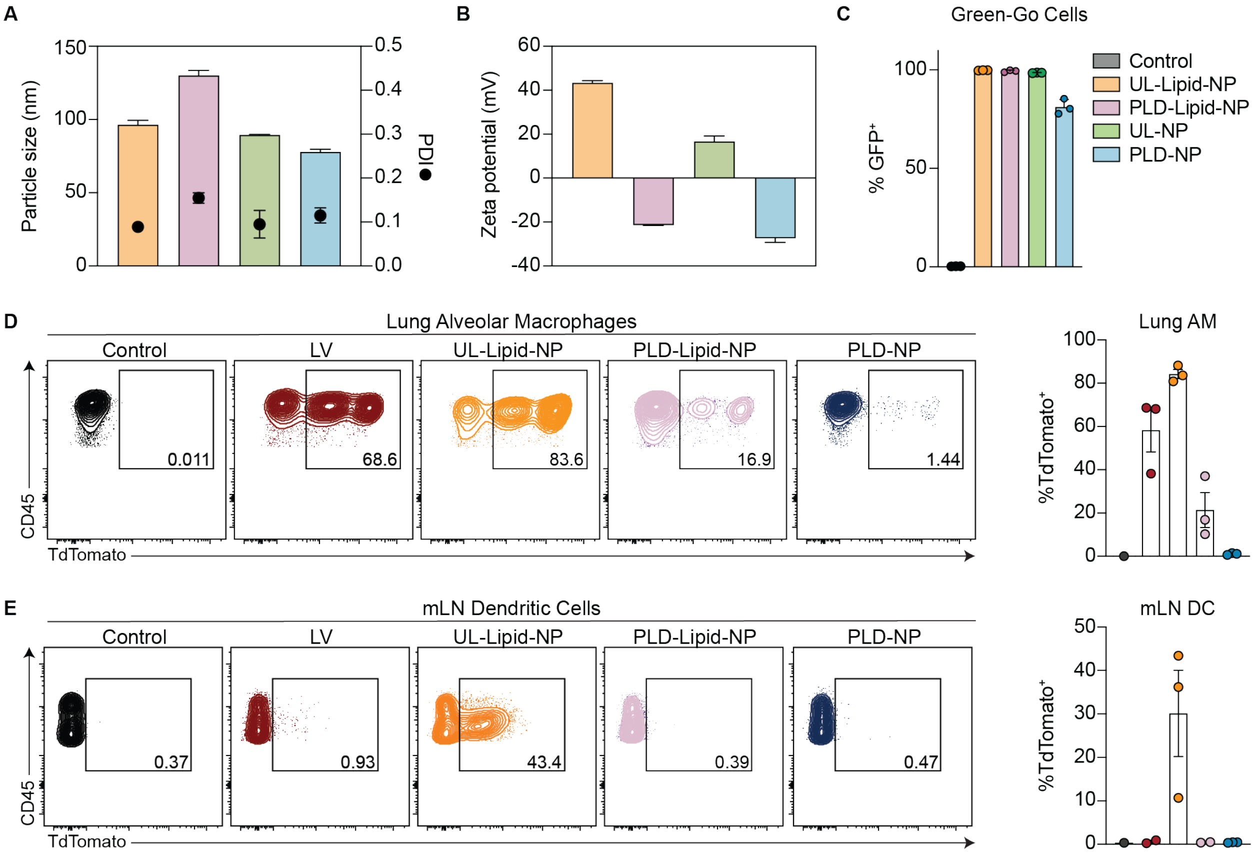


**Supplemental Figure 7.** A-B) Hydrodynamic diameter, PDI, and zeta potential of UL-Lipid-NP, PLD-Lipid-NP, UL-NP, and PLD-NP. C) Percent of GFP$^{+}$ Green-Go cells 72hr after transfection, n=3. D-E) Representative flow plots (left) and quantification (right) of TdTomato$^{+}$ AM (F) and DCs (G), 7 days after IT administration in T mice n=3.

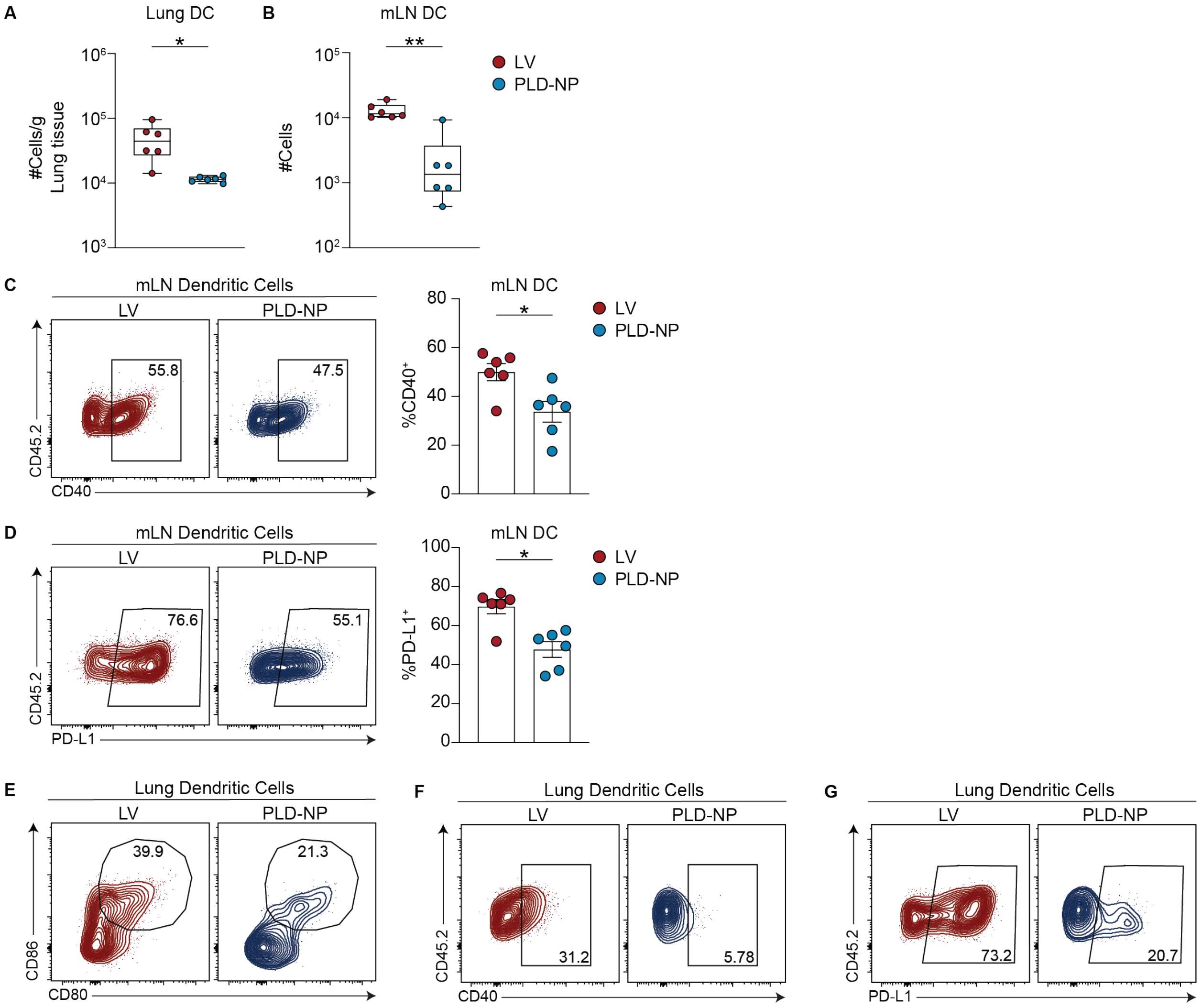


**Supplemental Figure 8.** A-B) Number of DC in the lung (A) and mLN (B) 7 days post I.T. injection, n=6. C-D) Representative flow plots (left) and quantification (right) of CD40 expression (C), and PD-L1 expression (D) on mLN DCs, n=6. For A-D, P-values calculated using Mann-Whitney U test. E-G) Representative flow plots of lung DCs expressing CD80 and CD86 (E), CD40 (F), and PD-L1 (G).

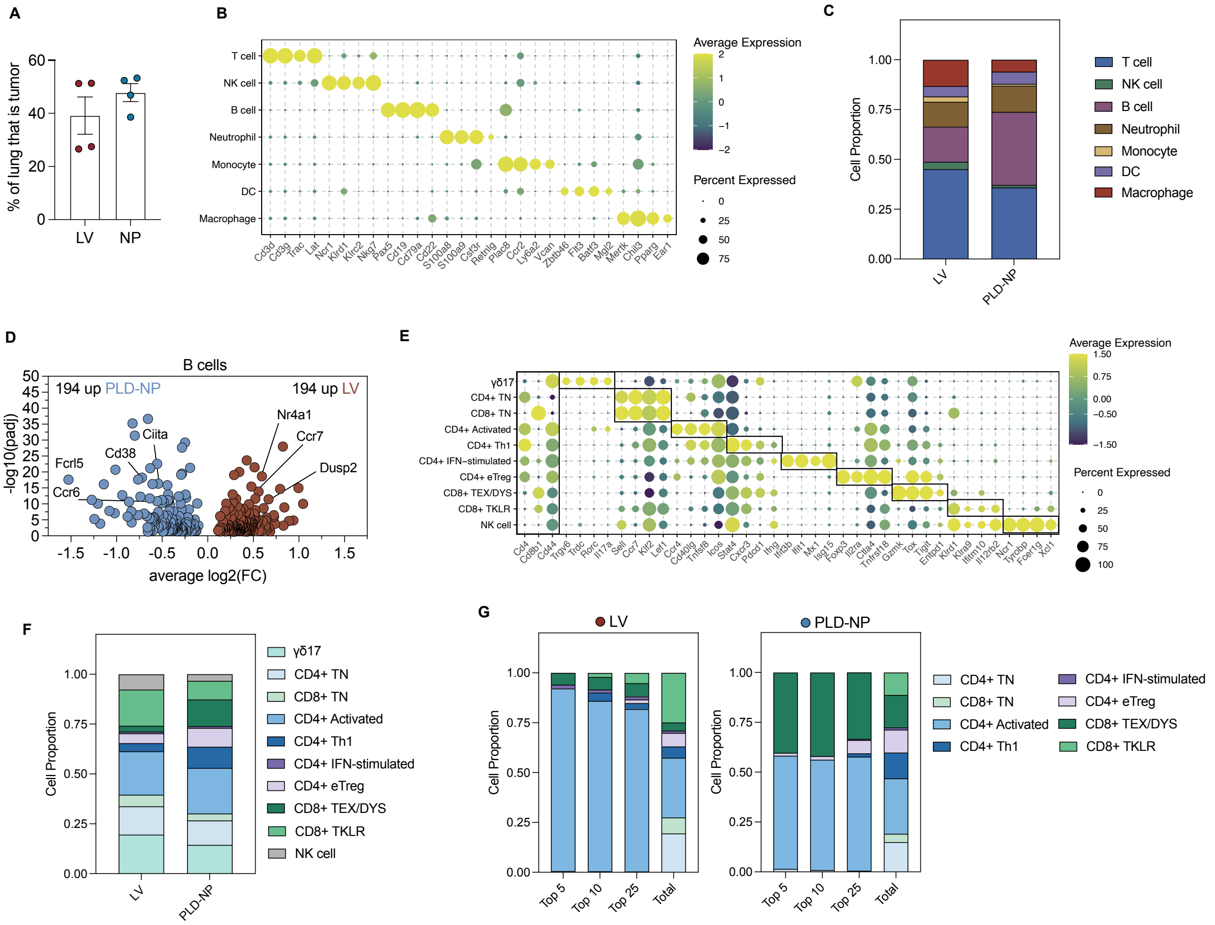

**Supplemental Figure 9**. (A) Tumor burden at 13 weeks post tumor initiation in KP lsl-SIY mice. (B)Dot plot of immune cell types and selected genes for each population. (C) Stacked bar plot of immune cell type composition per delivery method, expressed as proportion per cell type. (D) DGE between B cells from LV- vs. PLD-NP-induced tumors. (E) Dot plot of 10 T cell and NK cell phenotypes and selected genes characterizing each population. Cd4, Cd8b1, and Cd44 expression is provided on the left-hand side of the plot. (F) Stacked bar plot of T and NK cell phenotype composition per delivery method. (G) Stacked bar plot of T cell phenotype composition for top 5, 10, ad 25 abundant clonotypes per delivery method. The composition for all T cells (Total) is provided for comparison.
